# Concerted cutting by Spo11 illuminates DNA break mechanisms and initiates gap repair during meiosis

**DOI:** 10.1101/2019.12.18.881268

**Authors:** Dominic Johnson, Margaret Crawford, Tim Cooper, Corentin Claeys Bouuaert, Scott Keeney, Bertrand Llorente, Valerie Garcia, Matthew J. Neale

## Abstract

Genetic variation arises during meiosis via repair of DNA double-strand breaks (DSBs) created by the topoisomerase-like Spo11 protein. These DSBs are thought to always occur sparsely across the genome, with isolated DSBs generating discrete recombination events. We challenge this view, demonstrating that hyper-localised coincident DSBs frequently form within hotspots in both *S. cerevisiae* and mouse—a process suppressed by the DNA damage response kinase Tel1/ATM. Remarkably, the distances separating coincident DSBs vary with ∼10.5 bp periodicity, invoking a model where adjacent Spo11 molecules have a fixed orientation relative to the DNA helix. Deep sequencing of meiotic progeny identifies recombination scars consistent with gap repair initiated by adjacent DSBs. Our results revise current thinking about how genetic recombination initiates, reviving original concepts of meiotic recombination as double-strand gap repair.

## Results

### Mre11-independent formation of Spo11-oligonucleotide complexes

Despite more than 35 years since DSBs were proposed to initiate meiotic recombination^1^ and more than 20 years since identification of Spo11 as the DNA-cleaving protein^2,3^, it remains poorly understood how Spo11 engages its DNA substrate and how Spo11 DSBs are repaired. To make inroads, we examined the properties of Spo11-generated DSBs.

During DNA cleavage, Spo11 becomes covalently attached to each DNA end via a 5′ phosphotyrosyl link^2,4–6^ (**Fig. 1a, *left***). Spo11 is removed by the Mre11 nuclease^2,7–10^, generating Spo11 oligonucleotides (Spo11 oligos) (**Fig. 1a,b**) that enable Spo11 activity to be mapped within yeasts^11–13^, plants^14^ and mammals^15^. When separated by gel electrophoresis, deproteinised Spo11 oligos from *S. cerevisiae* form two prominent populations^9^ of ∼8-12 nt and ∼25-35 nt (**Fig. 1c**), the significance of which has so far remained unclear.

**Figure 1.**
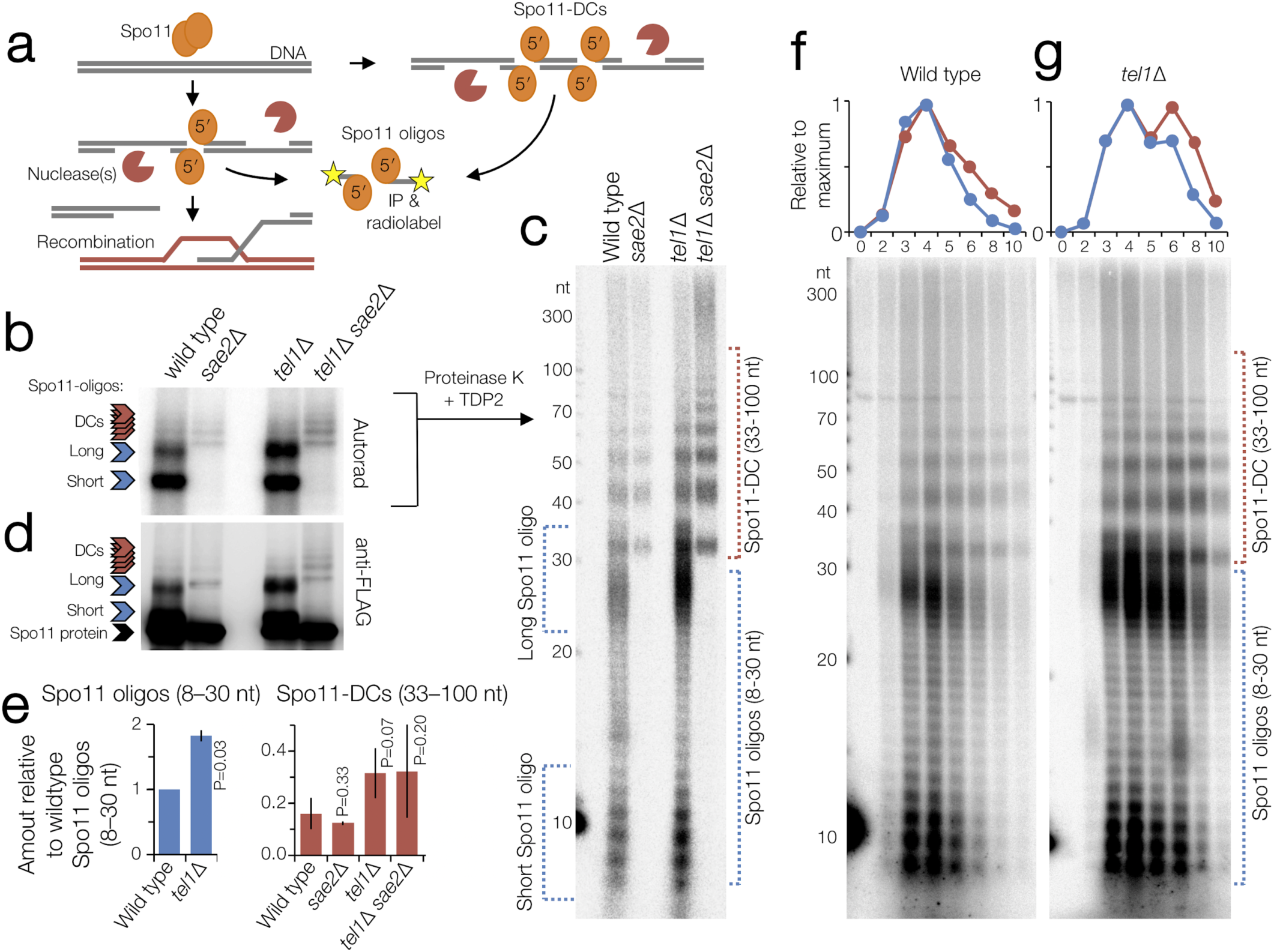
Spo11-oligo and Spo11-DC formation. **a**, Spo11 DSBs are nucleolytically processed to generate canonical covalent Spo11-oligo complexes. Adjacent Spo11 DSBs (“double cutting”, DC) can create a distinct class of Spo11-oligo complexes independently of nucleolytic processing. **b-g**, Detection of immunoprecipitated (IP) Spo11-oligo and Spo11-DC products by 8% SDS-PAGE (**b, d**) or 19% denaturing PAGE following treatment with proteinase K and TDP2 (**c, f-g**). Spo11 oligos and 10 nt marker standards were radiolabeled with chain-terminating [α-^32^P]-3′-dATP using terminal transferase (**b-c, f-g**). Total Spo11 detected via anti-FLAG western blotting (**d**). **e**, Quantification of signals (as in **c**) relative to wild type Spo11 oligos 8–30 nt in size (n=2; mean ± range is indicated; P=paired T test). **f-g**, Analysis of Spo11-oligo and Spo11-DC intermediates at hourly timepoints during meiotic prophase.

Advances in the sensitivity of Spo11-oligo analysis reveal additional higher molecular weight signals detectable via either radiolabelling of the 3′ OH group^16^ (**Fig. 1b,c**) or western blotting (**Fig. 1d**). The discrete banding of these Spo11 oligos is distinct from the heterogenous smear generated when Mre11 exonucleolytic activity is impaired^10^, and is instead reminiscent of molecules observed in mouse *Atm*^-/-^ mutants^17^ and in yeast lacking the *Atm* orthologue *TEL1*^16^.

We hypothesised that these longer Spo11 oligos could arise independently of Mre11 by Spo11 cleaving DNA at adjacent locations^16,18^ (**Fig. 1a, *right***). Moreover, such Spo11 double-cuts (Spo11-DCs) might be increased in *Atm*^*-/*-^ mutants, wherein DSB frequency increases ∼10-fold^17^. Consistent with this hypothesis, *sae2* mutants retained the ladder of higher molecular weight Spo11 oligos but not, as expected, oligos shorter than ∼30 nt (**Fig. 1b-d**). Sae2 is an essential activator of the Mre11 nuclease^19^.

High resolution gel analysis demonstrated that the ladder has a ∼10 nt periodicity—matching the helical pitch of B-form DNA—from ∼33 nt to >100 nt (**Fig. 1c, S1a,b**). The ladder was also visible when *SAE2* is present (**Fig. 1b-d**), and made up ∼14% of total Spo11 oligos in wild type. As with canonical (shorter) Spo11 oligos, the ladder was increased ∼2-fold in the absence of *TEL1* (**Fig. 1e**). Temporal analysis in wild type and *tel1*Δ revealed Spo11-DCs to arise concomitantly with Spo11 oligos, but to have longer apparent lifespan, suggesting greater stability (**Fig. 1f,g**).

Importantly, the ladder requires Spo11 catalytic activity (**Fig. S1c**), and is present in *mre11* and *rad50* separation-of-function mutants (**Fig. S1d**) that, like *sae2*Δ, cannot nucleolytically remove Spo11^8–10,20^. From these analyses we conclude that Spo11 often cleaves DNA at adjacent positions within both wild type and resection-deficient cells. If this idea is correct, we reasoned that the average size of Spo11-DCs should be increased in a strain where catalytically inactive (but DNA-binding competent) Spo11 is expressed alongside wild-type Spo11, because this should result in a greater average spacing between adjacent Spo11 complexes that are able to cut the DNA. This prediction was met (**Fig. S1e**) suggesting that inactive Spo11 complexes can bind DNA stably and act as competitive inhibitors of adjacent catalysis.

### Spo11-DCs are detected within whole-genome Spo11-oligo libraries

We reasoned that because Spo11-DC molecules are detected in the presence of *SAE2*, they may also be detected within the deep-sequencing Spo11-oligo libraries from wild type and *tel1*Δ strains^16^. To test this idea, we remapped the raw sequence reads using paired-end alignment, generating a length distribution of Spo11 oligos (**Fig. 2a**). In close agreement with the physical gel analyses, we detected not just the prominent class of previously characterised Spo11 oligos (20-40 nt), but also peaks at ∼43, ∼53, ∼63, and ∼73 nt (**Fig. 2a**). Moreover, a shoulder on the main peak was apparent at ∼33 nt in wild type and was more prominent in *tel1*Δ. Approximately 25% of the mapped reads are >40 nt in length. Based on our physical analysis, a proportion of these are likely to be Spo11-DCs.

**Figure 2.**
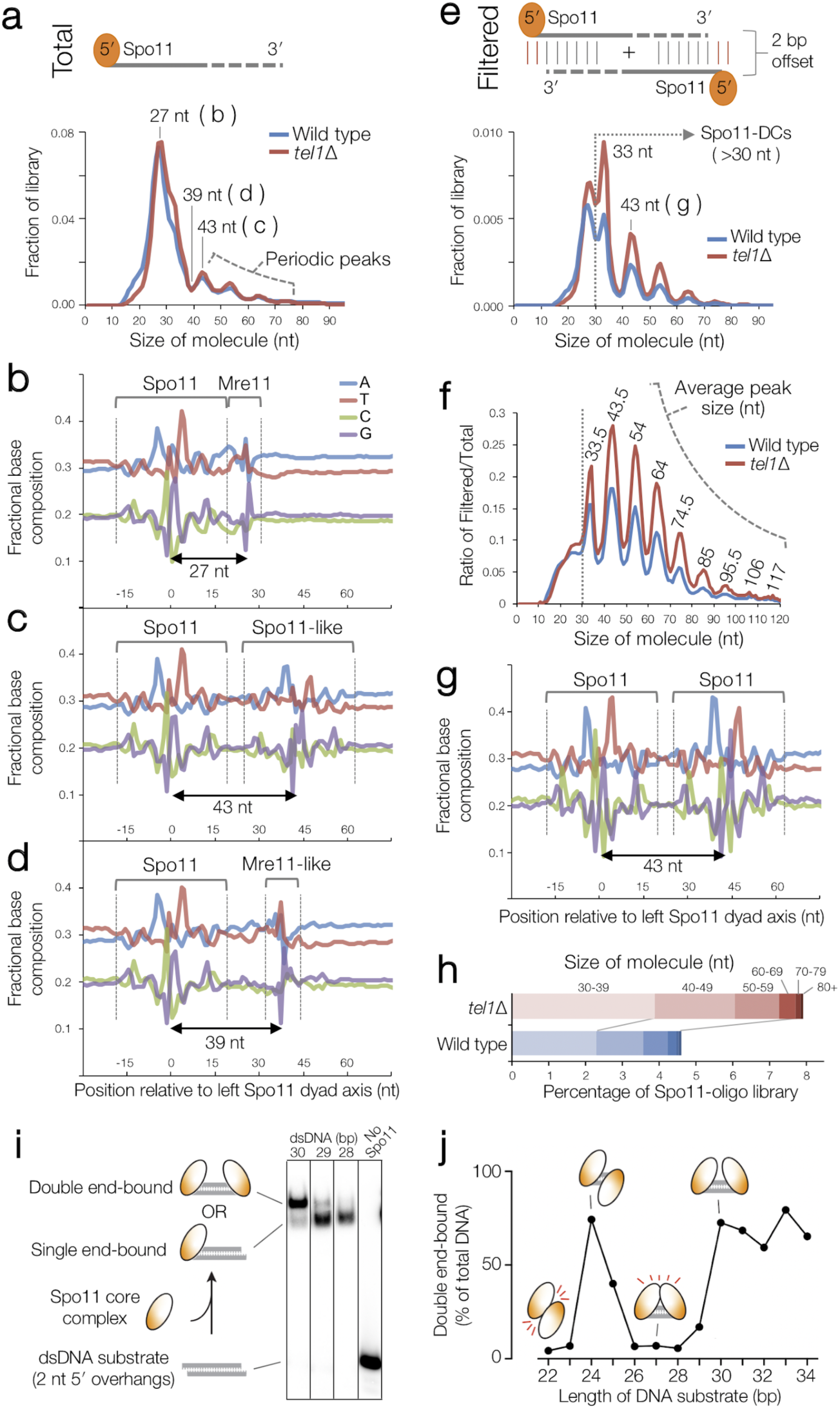
Identification of Spo11-DC within whole-genome Spo11-oligo libraries. Spo11-oligos^16^ isolated from the indicated strains were remapped using paired-end Bowtie2 alignment. **a**, Size distribution of all Spo11-oligos reveal periodic peaks >40 bp. **b-d**, Nucleotide composition of Spo11 oligos of the indicated size was computed for each base position, revealing Spo11 signature at both ends (**c**), or Spo11 signature at 5′ end plus Mre11 signature at 3′ end (**b, d**). **e**, Size distribution of the Spo11-oligo library after filtering to retain only those molecules that are part of a complementary pair with the expected 2 bp 5′ overhang created by Spo11 cleavage at both ends (Spo11-DCs). **f**, Ratio of filtered to total Spo11 oligos as a function of molecule length. **g**, Nucleotide composition of the 43-nt filtered molecules. **h**, Percentage of total Spo11-oligo library in the filtered (Spo11-DC) set for indicated size ranges. **i**, *In vitro* double-end binding assay. Spo11 associated with partners Rec102, Rec104 and Ski8 (‘core complex’) was incubated with short double-stranded DNA substrates with 2-nt 5′ overhangs, thereby mimicking the product of Spo11-mediated cleavage. The core complex binds such DNA ends with high affinity (Claeys Bouuaert *et al*., in preparation). **j**, The formation of double-end bound complexes was quantified with DNA substrates of different lengths. Double-end binding was efficient when the substrates were 25, 24 bp, and 30 bp or longer (**Fig. S2e**).

On average, Spo11 displays a characteristic DNA sequence bias spanning ±15 nt around the cleavage site, including preferred cleavage 3′ to a C nucleotide and flanking A/T skews^11,21^ (**Fig. 2b**). We reasoned that if the molecules ascribed to Spo11-DCs indeed arise from two coincident DSBs, the Spo11 oligos should have this sequence bias at both termini and not just the 5′ end.

To test this prediction, we stratified by Spo11-oligo length and assessed the base composition around the 3′ end of each molecule (**Fig. 2b-d**). As expected, 27-nt molecules within the major peak displayed a 3′ signature unlike Spo11 (**Fig. 2b**), attributed to sequence biases of Mre11 nuclease^11^. By contrast, 43-nt Spo11 oligos revealed a clear echo of the Spo11 sequence bias at their 3′ ends (**Fig. 2c**). Importantly, a Spo11-like bias was largely absent when assessing one of the troughs in the length distribution profile, 39 nt (**Fig. 2d**).

Next, we filtered the sequence reads to include only overlapping forward-strand (Franklin) and complementary-strand (Crick) read pairs with 5′ and 3′ ends separated by 2 bp (the overhang created by Spo11 cleavage^4,11^)—with the expectation that this will enrich for Spo11-DC molecules (**Fig. 2e**). Remarkably, this exercise enhanced the ∼10 nt periodicity even though the filter did not include an explicit constraint on read length (**Fig. 2e-f**). The Spo11-like sequence bias was also enhanced at the 3′ ends (**Fig. 2g**). These results strongly support the conclusion that the periodic Spo11-oligo lengths detected both within sequencing libraries (**Fig. 2e-f**) and physically (**Fig. 1**), are indeed mostly Spo11-DCs. Similar sequence biases were observed when selecting other peaks in the filtered distribution—including an otherwise undetectable peak at ∼33 bp (**Fig. 2e; Fig. S2a-d**).

Spo11-DCs (defined as filtered Spo11 oligos >30 nt; **Fig. 2e**) make up ∼4.6% and 7.9% of the total Spo11-oligo pool in wild type and *tel1*Δ strains, respectively (**Fig. 2h**). Presumably, these values are lower than our gel-based estimates due to size selection during library preparation and to the stringency of the filtering.

We hypothesized that the absence of Spo11-DCs <30 bp in length (**Fig. 1c** and **Fig. 2e,f**) is due to the inability of two DSB-forming complexes to assemble on DNA in such close proximity. This idea is consistent with the size of the region showing biased base composition (likely attributable to preferences in protein-DNA binding interactions) ±15 bp around each DSB end^11^ (e.g. **Fig. 2b**).

To directly test this idea, we examined recombinant Spo11-Rec102-Rec1104-Ski8 complexes, which can bind tightly and noncovalently to dsDNA ends in vitro (Claeys Bouuaert & Keeney, in preparation). Spo11 complexes were incubated with dsDNA fragments of varying length and assessed for their ability to bind one versus both ends (**Fig. 2i** and **Fig. S2e**). Remarkably, double-end binding was efficient for DNA molecules that were 30–34 bp or 24–25 bp, but not sizes in between (**Fig. 2j** and **Fig. S2e**). We interpret this result to mean that adjacent Spo11 complexes clash sterically with one another at distances below ∼30 bp if they are oriented in the same direction, but this clash can be alleviated by a relative rotation of 180° (half a helical turn) between the two Spo11 complexes (**Fig. 2j**).

Because molecules shorter than 30 nt are not detected in our physical and filtered Spo11-oligo analyses, we propose that adjacent Spo11 complexes capable of making double cuts must have interacted with DNA from the same orientation. Collectively, our data support the view that Spo11 can catalyse adjacent DSBs on the same DNA molecule; that Spo11-DCs arise much more frequently than current thinking had supposed; and that DSB centres in Spo11-DCs are separated by distances that increment in ∼10-bp steps from a minimum of ∼33 bp (approximately three helical turns of B-form DNA).

### Mapping Spo11 double-cuts within DSB hotspots

In *S. cerevisiae* and mouse, Spo11 preferentially cleaves DNA within regions of low nucleosome occupancy, generating a punctate map of preferred DSB ‘hotspots’ distributed non-randomly across the genome^11,15^. To determine the relationship between Spo11-DSB hotspots and Spo11-DCs, we generated comparative genome-wide maps of both, in which Spo11-DCs are represented as frequency-weighted arcs that link the 5′ ends of overlapping Franklin- and Crick-strand filtered reads (**Fig. 3a-c** and **Fig. S3a**).

**Figure 3.**
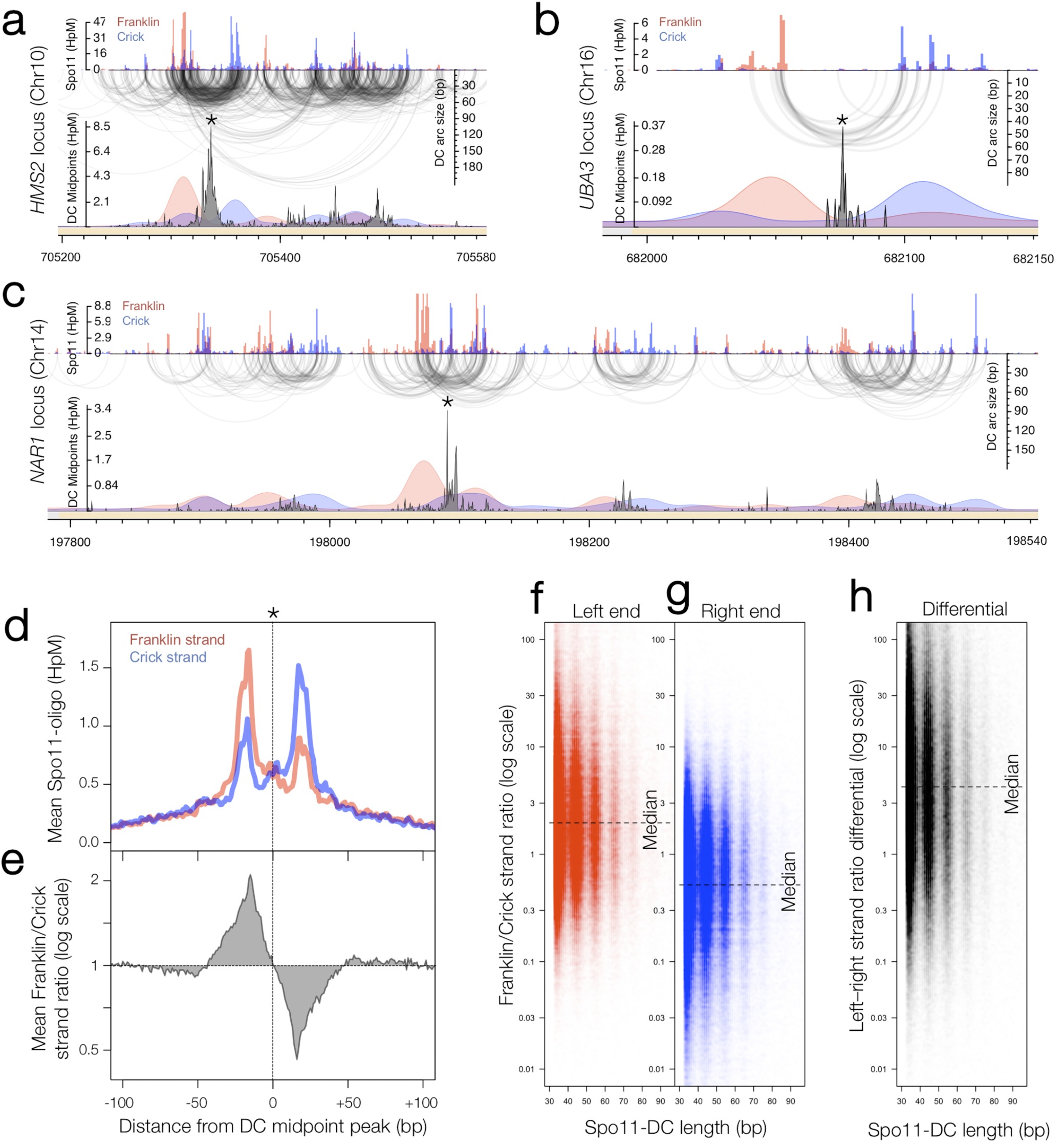
Fine scale analysis of Spo11-DCs within DSB hotspots. **a-c**, Arc diagram of Spo11-DCs mapped within three representative hotspots: strong (**a**), narrow (**b**), wide/complex (**c**). Top panel, unfiltered strand-specific Spo11 oligos (Franklin, red; Crick, blue; HpM, hits per million mapped reads). Arcs link 5′ ends of each Spo11-DC (i.e., filtered Spo11 oligos, **Fig. 3a)**. Lower panel, smoothed unfiltered strand-specific Spo11-oligos, overlaid with frequency histogram of Spo11-DC midpoints (grey). Note the spatial association of Spo11-DC sub-domains with Franklin-Crick strand disparity. **d-h**, Strand disparity at Spo11-oligo 5′ ends is a genome-wide feature of Spo11-DCs. Average strand-specific Spo11-oligo signal (**d**), and strand ratio (**e**), centred upon the strongest Spo11-DC midpoint (asterisks in **a-c**) within every annotated DSB hotspot (3910 loci). Strand ratio (Franklin/Crick for total Spo11-oligo HpM) was computed at the left and right 5′ end of every unique Spo11-DC molecule (**Fig. S3b**), stratified by length (**f-g**). Strand ratio differential (**h**) indicates the fold difference in the ratios when comparing the left and right 5′ ends of each Spo11-DC molecule (**Fig. S3b**).

Spo11-DCs are distributed non-uniformly across hotspots, with evidence of subdomains identifiable as peaks in Spo11-DC maps (**Fig. 3a** and **Fig. S4a**). Narrower hotspots display a simple pattern with centrally focussed Spo11-DC formation (**Fig. 3b** and **Fig. S4b**), whereas very wide hotspots contain multiple discrete Spo11-DC zones **(Fig. 3c** and **Fig. S4c**).

Overall, nearly all (∼95%) Spo11-DCs map within hotspots (**Fig. S5a**), more so than total Spo11 oligos (∼86%), suggesting that Spo11-DCs are more prevalent where Spo11 activity is strongest. Although the proportion of Spo11-DCs within each hotspot varied widely from <0.1% to >10% of the Spo11-oligo signal, ∼86% of hotspots displayed a Spo11-DC fraction of at least 1%, and ∼18% of hotspots displayed Spo11-DC fractions over 5% (**Fig. S5b**). In *tel1*Δ, these fractions increased to ∼94% and ∼47%, respectively (**Fig. S5b**), consistent with the median frequency of Spo11-DCs per hotspot being ∼1.7-fold greater (**Fig. S5b**; P<0.0001, Kruskal-Wallis H-test). Whilst Spo11-DC frequency correlated positively with total Spo11-oligo counts (**Fig. S5c**), the relationship was nonlinear, such that Spo11-DCs were observed disproportionately more frequently within the strongest hotspots (**Fig. S5d**). Finally, although Spo11-DCs were globally more frequent in *tel1*Δ compared to wild type, this relationship was not uniform across all hotspots (**Fig. S4d** and **Fig. S5e**).

### Spo11-DCs arise within regions of Spo11-oligo strand disparity

Every DSB is expected to generate two Spo11 oligos, one each mapping to the Franklin and Crick strands of the genome, but many individual DSB sites display strong disparity between strands^11^. Because Spo11 oligos <15 nt are not mapped, this disparity was thought to reflect differences in the relative numbers of longer and shorter canonical Spo11 oligos^9,11^. However, the molecular explanation for strand disparity at some but not all cleavage sites has been unclear.

Critically, we found an unanticipated alternating pattern of strand-biased Spo11-oligo enrichment across hotspots that was spatially associated with Spo11-DC subdomains (**Fig. 3a-c**). Specifically, the left flanks of Spo11-DC peaks appeared to be enriched for Franklin-strand hits, whereas the right flanks were enriched for Crick-strand hits (**Fig. 3a-c** and **Fig. S4a-d**). This pattern was most pronounced at narrow, low frequency hotspots (**Fig. 3b** and **Fig. S4b**) and was retained upon *TEL1* deletion (**Fig. S4d**).

To investigate the generality of these observations, we assessed strand-specific Spo11-oligo signal when averaged around the centres of the strongest Spo11-DC peaks in all ∼3900 annotated DSB hotspots (**Fig. 3d**). Mapped 5′ ends of bulk Spo11 oligos displayed peaks ±20 bp from Spo11-DC centres and were associated with ∼2-fold skews towards Franklin-strand hits on the left and Crick-strand hits on the right (**Fig. 3e**). A similar disparity in local Spo11 oligos was observed at the left (**Fig. 3f**) and right (**Fig. 3g**) ends of individual Spo11-DCs regardless of their size, resulting in a ∼4-fold differential towards Franklin on the left and Crick on the right (**Fig. 3h**), also seen upon *TEL1* deletion (**Fig. S6a-d**). Importantly, no net disparity was observed within the bulk Spo11-oligo population (**Fig. S6e-f**), indicating that these strand-specific patterns are a special feature of genomic sites that generate Spo11-DCs. We conclude that our observations are not unique to specific hotspots, nor to specific mutants, but are, instead, a general feature of the meiotic recombination process.

### Spo11-DCs lead to double-strand gap repair

The molecular data presented so far show that closely spaced DSBs can arise on the same DNA molecule. Because Spo11-DCs are recovered in a wild-type background, we infer that resection initiation is inefficient in between such coincident DSBs. Therefore, Spo11-DCs might be repaired as short double-strand gaps, leading to unidirectional transfer of genetic information from the uncut donor DNA to the gapped molecule (**Fig. 4a**).

**Figure 4.**
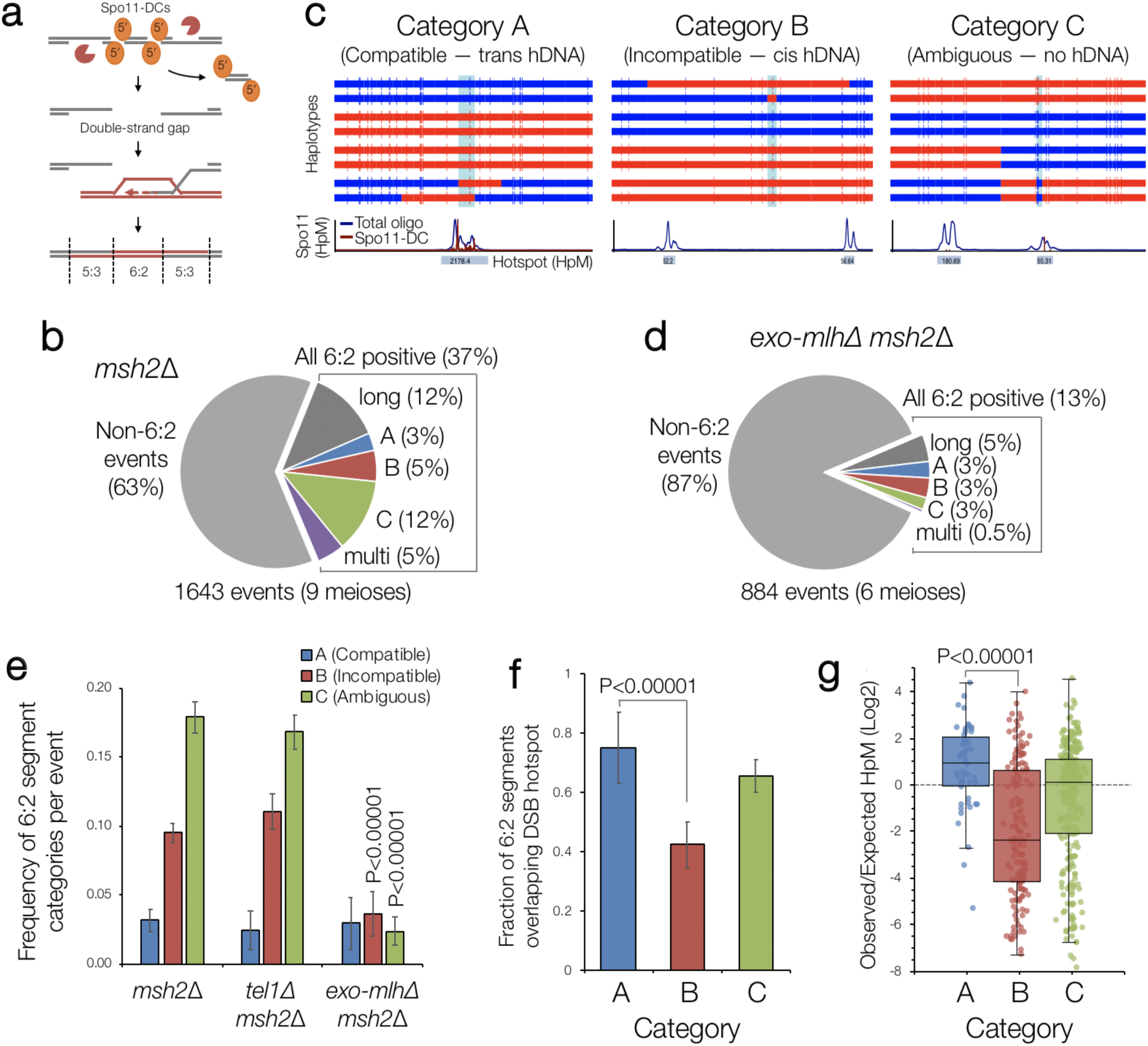
Identification of gap repair events during meiosis. **a**, Gap repair is predicted to generate segments of 6:2 marker segregation flanked by 5:3 segments in mismatch repair-deficient strains. **b-d**, Quantification of recombination event types in (**b**) control (*msh2*Δ), and (**d**) pooled *exo1*Δ *msh2*Δ, *mlh1*Δ *msh2*Δ, *mlh3*Δ *msh2*Δ (‘*exo-mlh*Δ *msh2*Δ’) strains based on representative categorisations presented in (**c**) based on the patterns of flanking heteroduplex DNA marker segregation (trans, cis, none). Upper panels in (**c**) are the genotype calls made at each SNP marker (vertical line). Adjacent segments of the same genotype are joined with horizontal bars (red or blue) to aid visualisation of patterns. Each horizontal bar is sequenced haplotype from one meiotic octad. 6:2 segments are indicated in pale blue. Lower panels are smoothed histograms of total Spo11-oligo (blue) and the filtered Spo11-DC signals (purple). **e**, Frequency of 6:2 segments per event was calculated for each category and genotype. P values indicate Z-test of proportions relative to *msh2*Δ. **f**, Fraction of 6:2 segments (*msh2*Δ) overlapping annotated hotspots. **g**, Log_2_ ratio of observed Spo11-oligo density within each 6:2 segment divided by the mean Spo11-oligo density within the entire event, in *msh2*Δ, for each category. P values indicate Z-test of proportions (**f**) and Kruskal-Wallis H-test (**g**).

To test this idea, we analysed meiotic recombination ‘scars’ in a hybrid yeast strain containing ∼65,000 nucleotide sequence polymorphisms^22,23^. To reduce analytical ambiguity from correction of mismatches in heteroduplex DNA, we deleted the mismatch repair factor *MSH2*^22,23^. In *msh2*Δ cells, recombination events initiated by a single DSB are expected to generate heteroduplex DNA, visible as segments of 5:3 marker segregation within a recombination scar^22^. By contrast, 6:2 marker segregation should be enriched within events initiated by gaps (**Fig. 4a**). As previously noted^22^, a large fraction (∼37%) of recombination events in *msh2*Δ cells contained at least one patch of 6:2 segregation (**Fig. 4b**). We filtered these events to include only those with 6:2 segments between 30 and 150 bp (∼25% of total events), consistent with the sizes of Spo11-DCs determined physically.

Next, because 6:2 patches can also be generated by mechanisms other than adjacent Spo11 DSBs^23^, we sorted events into three categories (A–C) based on the flanking patterns of heteroduplex DNA (**Fig. 4c** and **Fig. S7a**. See **Methods** for details). Category A events are compatible with gap repair because flanking heteroduplex DNA patterns are in trans orientation, whereas category B events are incompatible because the flanking heteroduplex DNA patterns are in cis orientation. Category C events lack the flanking heteroduplex DNA patterns necessary to assign the event.

Category A (compatible) accounts for ∼3% of all events, category B (incompatible) is ∼5% and category C (ambiguous) is ∼12% (**Fig. 4b** and **Fig. S7a-g**). Loss of Tel1 had no impact on the relative proportion of categories A–C (**Fig. S7a** and **Fig. S8a**), but increased the frequency of events with 6:2 segments >150 bp (**Fig. S8a**), consistent with a loss of DSB interference increasing the probability of coincident DSBs at adjacent hotspots^18^.

6:2 segments that do not arise from Spo11-DCs may arise from the DNA nicking action of Mlh1-3 and Exo1 during repair^23^. Importantly—and consistent with this idea—abrogation of these nicking activities led to a specific reduction in the frequency of category B and C events, whereas category A events were unaffected (**Fig. 4d** and **Fig. S7a**; P<0.0001 two-tailed Z-test).

We further reasoned that 6:2 segments arising from gap repair should correlate strongly with the location of DSB initiation, whereas nick translation during repair has the potential to arise away from the initiation site. Thus, to further test the validity of these categorisations, we computed both the frequency of 6:2 segments overlapping Spo11 hotspots (**Fig. 4f**), and the density of Spo11 oligos within the 6:2 segments (**Fig. 4g**) of all categories. By both metrics, Category A events displayed the greatest association with Spo11 activity (P<0.00001, two-tailed Z-test and Kruskal-Wallis H test, respectively). Collectively, these analyses support the view that category A events arise from gap repair.

### Spo11-DCs arise during mouse meiosis

DSB formation by Spo11 is evolutionarily conserved^24^. Notably, in the absence of ATM, mouse SPO11 oligos are more abundant and display a distinct size distribution, including a ladder of molecules larger than the canonical SPO11-oligo signal^17^ similar to those in *S. cerevisiae*.

To investigate whether SPO11-DC formation occurs in mammals, we explored SPO11-oligo sequences from mouse spermatocytes^15^. Mouse SPO11-oligo size distribution is distinct between wild type and *Atm*^*-/-*^, with prominent populations at ∼10–20 nt and ∼25–30 nt in the wild type, and larger molecules (25–60 nt) more abundant in *Atm*^-/-^ **(Fig. 5a**). Filtering to retain only overlapping SPO11 oligos, whilst less efficient than in *S. cerevisiae*, disproportionately retained putative mouse SPO11-DCs in the 30–60 nt range in *Atm*^*-/-*^ (**Fig. 5b-c**). Moreover, subtle periodic peaks of enrichment at ∼32 nt, 43 nt, and 53 nt emerged, reminiscent of *S. cerevisiae* Spo11-DC sizes (**Fig. 5c**).

**Figure 5.**
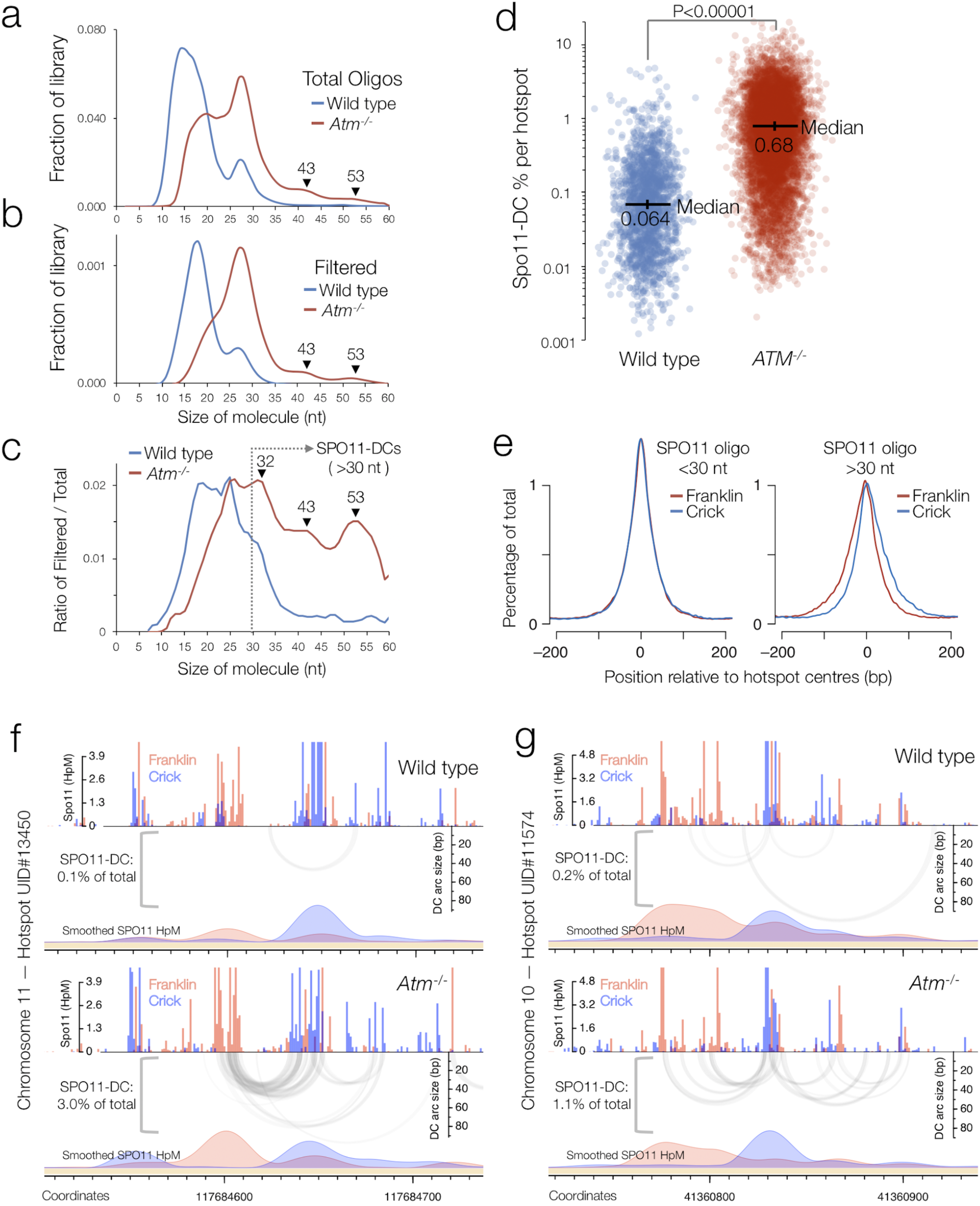
Fine-scale analysis of SPO11-DCs within mouse DSB hotspots. Mouse SPO11-oligo libraries^15^ were remapped using paired-end Bowtie2 alignment. **a-c**, SPO11-oligo length distribution of the entire (**a**) and reciprocally filtered (**b**) library enriches peaks in *Atm*^-/-^(**c**) that display a ∼10 bp periodicity. As for *S. cerevisiae*, SPO11-DCs are defined as filtered molecules >30 nt in length. **d**, Percentage of total SPO11 oligos that are SPO11-DCs, plotted for every hotspot. P value indicates Kruskal-Wallis H-test. **e**, Total *Atm*^-/-^ SPO11 oligos were filtered into two size classes then aggregated around ∼21,000 hotspot centres revealing a strand-specific disparity for Spo11-oligos >30 nt. **f-g**, Representative arc diagrams of SPO11-DCs (grey-scale frequency-weighted arcs) in wild-type and *Atm*^*-/-*^ relative to total strand-specific SPO11 oligos (upper, raw; lower, smoothed; red, Franklin; blue, Crick). Percentage of total SPO11 oligos that are SPO11-DCs is indicated.

Filtered SPO11 oligos were approximately 10-fold more enriched within annotated hotspots in *Atm*^*-/-*^ than the control (**Fig. 5d**; P<0.0001, Kruskal-Wallis H test). Interestingly, bulk SPO11-oligo signals >30 nt in length were offset from hotspot centres in a strand-specific manner reminiscent of the left-right Franklin-Crick disparity observed at *S. cerevisiae* hotspots (**Fig. 5e**). Arc maps revealed alternating domains of strand disparity associated with locations of filtered SPO11 oligos (**Fig. 5f-g** and **Fig. S9**), similar to *S. cerevisiae* (**Fig. 3**). These observations suggest that Spo11-DC formation is evolutionarily conserved.

## Discussion

Although double-strand gap repair assays were instrumental in establishing the DSB repair model more than 35 years ago^1^, direct Spo11-dependent gap formation in vivo was not anticipated, nor does it feature within current models. Indeed, despite the DNA interaction surface of Spo11 likely being <30 bp^11,25^, it has been generally assumed that only a single DSB is created within any given hotspot at a time even though most meiotic hotspots are many times wider^11,15^ (100-1000 bp).

Challenging this idea, we show that adjacent Spo11 dimers can cleave coincidentally at close proximity within hotspots, and we provide evidence that Spo11-DCs yield double-strand gaps that cause mismatch repair-independent unidirectional transfer of genetic information. Our conclusions agree with and substantially extend interpretations of segregation patterns in mismatch repair-defective strains^22^, and Spo11-oligo sizes in *tel1* mutants^16^.

The ladder of yeast Spo11-DCs represents ∼14% of total Spo11 oligos. A Spo11-DC should generate four Spo11-oligo complexes instead of the two oligos generated at a single Spo11 DSB. If we assume that there is at most only one Spo11-DC per recombination initiation event, we estimate that ∼16% of events contain a Spo11-DC (**Methods**). The lower frequency estimated for gap repair in recombination data (∼3%) likely reflects uncertainty in ascribing recombination events to gap repair, inability to detect intersister recombination, and effects of limited polymorphism density (1 per ∼200 bp), which disproportionately reduces the chance of detecting gaps the shorter they are (**Fig. S7h**).

The ∼10-bp periodicity of Spo11-DC sizes is intriguing. Similar periodicity occurs for DNase I cleavage of DNA when wrapped around histones^26^. Within nucleosomal DNA, the periodicity arises due to only one face of the helix being solvent exposed, with the alternating major-minor groove pattern then repeating once per helical turn. Importantly, however, DSB hotspots have a largely open chromatin structure, and stable nucleosomes occlude Spo11 access to DNA in vivo^11^, similar to other topoisomerase family members^27^. Therefore, we consider it likely that something other than nucleosomes restricts Spo11-DC endpoints to these periodic positions.

We envision a mechanism wherein multiple Spo11 proteins create a platform that enables coordinated Spo11-DSB formation (**Fig. 6a**). In this model, periodic spacing arises from adjacent Spo11 molecules being restricted to accessing the same face of the DNA helix, with preferred cleavage opportunities arising only every helical turn (∼10.5 bp) because of the relative inflexibility of B-form DNA over these short distances. This model also explains why the minimum Spo11-DC size is ∼33 bp despite our DNA binding observations indicating that adjacent Spo11 complexes can come as close as 24 bp in vitro when not constrained to be in the same orientation relative to the DNA helical axis. Moreover, a multimeric DSB-forming ‘machine’ made up of many catalytic centres may explain the previously perplexing observation that *SPO11*/*spo11-Y135F* catalytic heterozygosity has little effect on overall DSB formation despite the expectation of negative dominance^25^.

**Figure 6.**
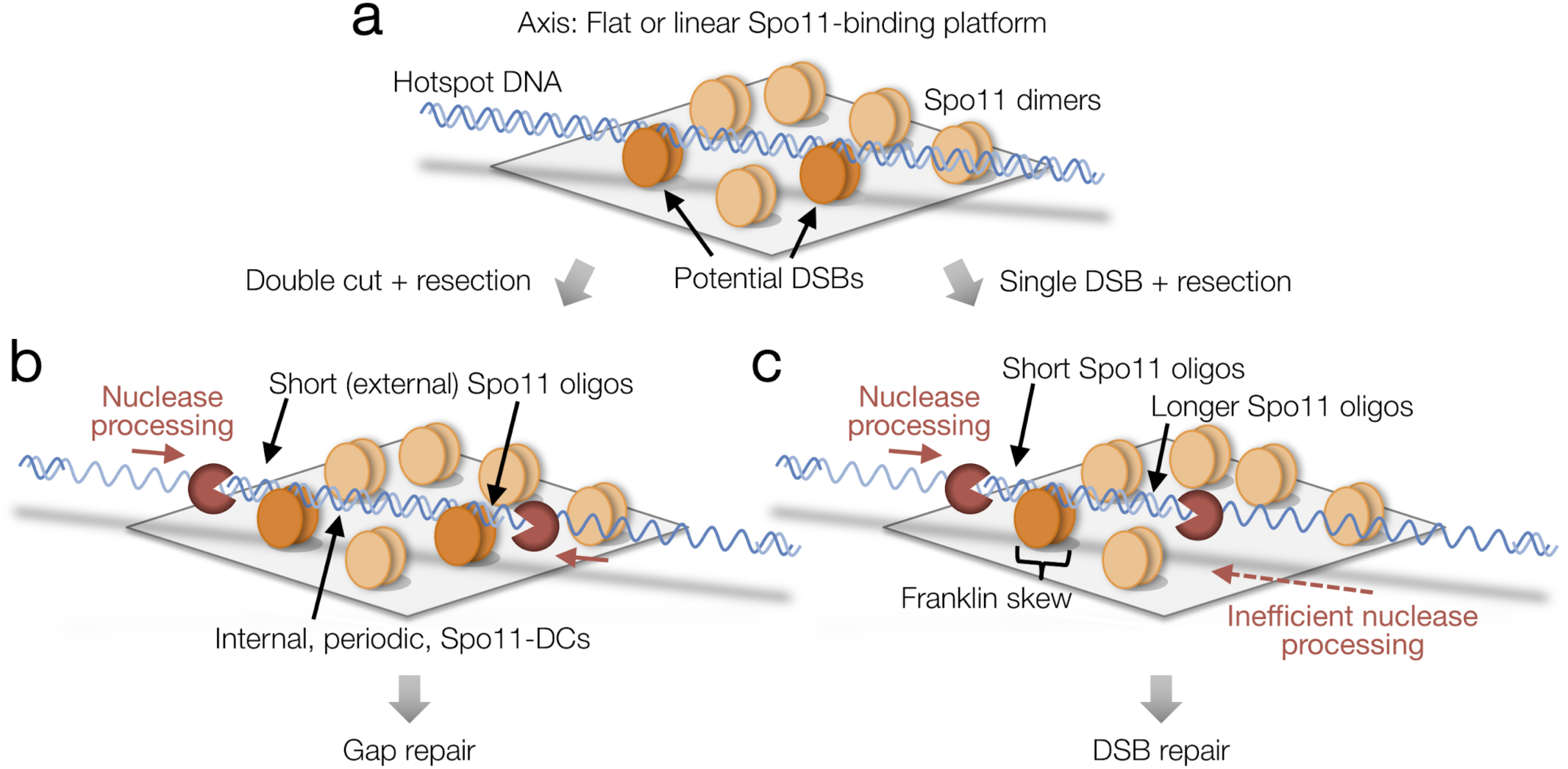
Model for Spo11-DC formation. **a**, Hotspot DNA interacts with multiple Spo11-dimers (pale orange ellipses) bound within a relatively flat surface. Single or coincident DSBs may form depending upon spatially-periodic favoured interactions between Spo11 and the repeating structure of the DNA helix (dark orange Spo11 dimers). **b**, Coincident DSB formation generates internal Spo11-DCs with lengths that are multiples of the helical pitch (∼10.5 bp). Resection initiates efficiently in the flanking regions generating short, external, Spo11-oligos, and inducing gap repair. **c**, Single DSBs are processed by nucleases on both sides, but resection proceeds less efficiently within the axis-associated DNA, generating a Spo11-oligo length asymmetry and consequent mapped strand disparity (see also **Fig. S9**).

In our model, the alternating strand disparity in Spo11-oligo libraries arises from the Spo11 platform selectively protecting a subset of DSB ends from the endonuclease and/or 3’→5’ exonuclease activities of Mre11^10^. For DSB ends within the platform, or facing inwards from the platform edges, this protection would yield Spo11-DCs as well as the canonical Spo11 oligos of larger sizes (**Fig. 6b-c** and **Fig. S3d**). In contrast, the outermost DSB ends would be less protected, yielding Spo11 oligos vulnerable to more extensive digestion by Mre11, in turn leading to their under-representation in Spo11-oligo maps that omit short oligos (**Fig. 6b-c** and **Fig. S3d**). Thus, we propose that the two prominent size classes of Spo11 oligo detected physically^9^ arise from asymmetric hotspot–axis interactions that generate differential sensitivity to nucleolytic degradation.

In broad terms, our model for a surface-bound Spo11 platform agrees with how Spo11 and its essential cofactors interact with the chromosomal axis as measured by ChIP^28–30^ and by immunofluorescent staining of spread chromosomes^31–33^, such that Spo11 would be organised alongside other pro-DSB factors in a two-dimensional protein array. Whilst we draw the axis in a planar form, our ideas do not exclude a model where the axis and DNA curve or writhe in concert with one another, and it is upon the exposed surface of such a structure that Spo11 catalysis occurs. However, the fact that we detect no major DNA sequence skews towards more flexible A or T bases in the centre of Spo11-DC fragments (**Fig. S2d**), disfavours the idea that DNA is subject to significant localised bending forces during Spo11-DC formation.

Tel1/ATM has an evolutionarily conserved function in negatively regulating DSB numbers^15–18,34–36^. In yeast, at hotspots ∼2 kb apart, loss of Tel1-mediated DSB interference increases coincident DSB formation ∼10-fold^18^. By contrast, *TEL1* deletion increases Spo11-DCs much less (∼2-fold; **Fig. 1b-e**), despite each Spo11-DC also being formed by a pair of adjacent DSBs. We interpret this difference to indicate that—unlike DSBs arising in two separate hotspots—the closely spaced DSBs that generate a Spo11-DC are less subject to Tel1-dependent negative regulation between one another. We envisage a mechanism where, in response to a DSB, Tel1 inactivates adjacent patches of assembled Spo11 in which DSB formation has yet to happen. By contrast, we propose that coincident DSBs often arise within a hotspot before the inhibitory effect of Tel1 can act—promoted by the high-density Spo11 array (**Fig. 6**).

Our exploration of mouse Spo11-oligo libraries supports the view that Spo11-DC formation within meiotic hotspots is evolutionary conserved—an interpretation consistent with recent molecular analysis of recombination in *Atm*^*-/-*^ mouse spermatocytes (A. Lukaszewicz, S. Keeney, and M. Jasin, unpublished). However, the frequency of SPO11-DCs increases much more in mouse *Atm*^*-/-*^ than it does in *tel1*Δ yeast (∼10-fold, **Fig. 5d** vs ∼2-fold, **Fig. 2h**). We interpret this difference to suggest that SPO11 double cutting in mouse is subject to direct inhibition by ATM. Such an effect may be a consequence of the more central role ATM plays within the generalised DNA damage response in mammals. Alternatively, or in addition, it may point to as-yet-uncharacterised differences between how SPO11-DSB formation is controlled within the two organisms.

For many years, a mechanistic understanding of the nature of Spo11-DSB formation has remained out of reach. The features of Spo11-oligo and Spo11-DC sizes and locations broaden understanding of the meiotic recombination pathway to include concerted DSB formation and gap repair as a frequent and evolutionarily conserved feature of meiotic recombination, and provide a tantalising glimpse into how the elusive biochemistry of Spo11 works in vivo.

## Acknowledgements

We thank Shintaro Yamada for help analysing the mouse Spo11-oligo datasets, Marie-Claude Marsolier-Kergoat for sharing Python scripts, Jesús Carballo and Michael Lichten for sharing *S. cerevisiae* strains containing relevant constructs (*tel1*Δ::hphNT2 and *sae2*Δ::kanMX6, respectively), and Rachal Allison for critical reading of the manuscript.

## Author contributions

M.J.N. and V.G conceived the project and prepared the manuscript. D.J., V.G, C.C.B., and M.J.N performed physical analysis of Spo11-DCs. M.C. prepared and analysed whole genome recombination maps with B.L. advising upon the mechanistic interpretation. T.C. and M.J.N mapped and analysed Spo11-oligo library data. C.C.B. and S.K. contributed protein biochemistry and provided critical mechanistic interpretations. All authors helped write the manuscript.

## Data availability

Raw *S. cerevisiae* and mouse Spo11-oligo FASTQ data were obtained from published archives GSE84896 and GSE84689 respectively. Nucleotide-resolution maps generated by paired-end Bowtie2 alignment are provided as supplementary files. The *S. cerevisiae* maps generated here were equally-mixed pools of the following biological samples: wild type, GSM2247761–GSM2247765; *tel1*Δ, GSM2247766–GSM2247769, GSM2354276, and GSM2354277). For mouse, maps used here were generated from the following biological samples: wild type, GSM2247728; *Atm*^-/-^, GSM2247731. FASTQ files used for mapping HR patterns in *S. cerevisiae* octads in *msh2*Δ and *tel1*Δ *msh2*Δ are deposited in the following NCBI SRA archives, PRJNA479661 and PRJNA480956, respectively.

## Methods

### Strains and culture methods

*S. cerevisiae* strains used in this study are listed in **Table S1**. Synchronous meiotic cultures were grown using standard methods. Briefly, YPD cultures (1% yeast extract, 2% peptone, 2% glucose) were diluted 100-fold into YPA (1% yeast extract, 2% peptone, 1% K-acetate) and grown vigorously for 14 h at 30°C. Cells were collected by centrifugation, washed once in water, resuspended in an equal volume of prewarmed 2% K-acetate containing diluted amino acid supplements, and shaken vigorously at 30°C. For mapping meiotic recombination patterns in hybrid octads SK1 and S288c haploid parents were mated for 8– 14 hours on YPD plates as described^37^. Cells were washed and incubated in sporulation media at 30°C with shaking, and tetrads were dissected after 72 hours. To generate octads, dissected spores were allowed to grow for 4-8 hours on YPD plates until they had completed the first post-meiotic division, after which mother and daughter cells were separated by microdissection, and allowed to grow for a further 48 hours. Spore clones were subsequently grown for 16 hours in liquid YPD prior to genomic DNA isolation using standard techniques^18^. Only octads generating eight viable progeny were used for genotyping by deep sequencing.

### Spo11-oligo and Spo11-DC physical analysis

Spo11-oligo complexes were isolated by immunoprecipitation using anti-FLAG antibody from 10 ml aliquots of sporulating cells harvested at the indicated timepoint(s), and labeled with alpha-32P cordycepin triphosphate using Terminal deoxynucleotidyl transferase (Fermentas) as described^38^, and separated by 7.5% SDS-PAGE, or treated with Proteinase K (Fisher) for 1 hour at 37°C prior to overnight precipitation in 90% ethanol at −80°C, then denatured in 1x formamide loading dye and separated on 19% denaturing urea/PAGE in 1x TBE. Where indicated, deproteinised samples were further treated with 300 nM recombinant mammalian TDP2 for 30 minutes at 30°C^38^ prior to electrophoresis, to remove residual phosphotyrosyl linked peptides, thereby enabling an accurate estimate of Spo11-oligo DNA length. Radioactive signals were collected on phosphor screens, scanned with a Fuji FLA5100 and quantified using ImageGauge software. For analysis of Spo11-oligo species by Western, SDS-PAGE gels were transferred to PVDF membrane, blocked with 5% non-fat dry milk (NFDM) / 1x TBST, and incubated with anti-FLAG-HRP at 1:1000 in 1% NFDM / 1xTBST. To estimate the fraction of events containing a Spo11-DC, we apply the simplifying assumption that, at most, only a single Spo11-DC arises per event such that each Spo11-DC makes four Spo11-oligos (two internal, two external) whereas each single Spo11 DSB makes only two oligos (two external). Applying these assumptions, the estimated fraction of events containing a Spo11-DC simplifies to the equation: Spo11 oligos_>30 nt_ / Spo11 oligos_<30 nt_

### Remapping of *S. cerevisiae* Spo11-oligo libraries

*S. cerevisiae* Spo11-oligo libraries^16^ were aligned to the Cer3H4L2 reference genome using Bowtie2, with identical GLOBAL and LOCAL mapping parameters: -X1000 --no-discordant --very-sensitive --mp 5,1 ----np 0. Cer3H4L2 is identical to the sacCer3 reference genome (R64-1-1), with the addition of two ectopic insertions: 1173 bp of hisG sequence inserted at the *LEU2* locus at position 91965, and 3037 bp of LEU2 sequence including 77 bp of associated unidentified bacterial sequence^39^ at position 65684. Before mapping, reads were trimmed to remove adapters and trailing G or C bases introduced during library preparation using Perl (OligoTrim.pl). Specifically, 5′ ends of Read1 were trimmed using Perl to remove the following sequences at the first 9 or 8 bases of each read: NNNNNCCCC (or NNNNNCCC if prior sequence not found). Read1 3′ ends were trimmed to truncate before any GGGGAGAT (or GGGAGAT if prior sequence not found) sequences should they be present. Similarly, Read2 5′ ends were trimmed to remove leading CCCC (or CCC if prior sequence not found) sequences, and 3′ ends truncated before GGGGNNNNNAGAT (or GGGNNNNNAGAT if prior sequence not found) sequences. In each case, the need to trim the NNNNN string arises from the use of custom barcoded adapters during library preparation^16^. The AGAT string is the reverse complement of the first 4 bp of the universal Illumina adapter. Following trimming and paired end alignment, the Read1 5′ base is expected to have a high probability of being a true Spo11-oligo 5′ end, and the terminal mapped base a true 3′ end. Nevertheless, some ambiguity is impossible to avoid due to inherent uncertainties in the number of terminal rG bases added during library preparation^16^. Resulting SAM files were processed via terminalMapper (https://github.com/Neale-Lab) using the ‘DOUBLE’ setting, generating 1 bp resolution histogram files of 5′ Spo11 oligo ends mapping to either Franklin or Crick strands of the genome. Additional “CoordinateAB” files report frequencies, strand, and position of molecules with unique 5′ and 3′ ends, enabling filtering for overlapping pairs. In these files, the 3′ reported is 2 nt more distal than the actual 3′ end such that it corresponds to any putative 5′ end on the complementary strand. As such, and because the AB coordinates listed include the first and last mapped base, the raw oligo lengths are 1 nt shorter than than obtained by subtracting the B coordinate from A.

### Remapping of mouse Spo11-oligo libraries

Mouse *Atm*^-/-^ SPO11-oligo libraries^15^, were trimmed in a similar way. Specifically, 5′ ends of Read1 were trimmed using Perl to remove the following sequences at the first 9 or 8 bases of each read: NNNNNCCCC (or NNNNNCCC if prior sequence not found). Read1 3′ ends were trimmed to truncate before any GGGG (or GGG if prior sequence not found) sequences should they be present. Similarly, Read2 5′ ends were trimmed to remove leading CCCC (or CCC if prior sequence not found) sequences, and 3′ ends truncated before GGGG (or GGG if prior sequence not found) sequences. Resulting SAM files were processed with terminalMapper as above, but additionally processed to expand the called 5′ and 3′ ends by ±1 bp in a manner that averages the frequency of reads mapped at a specific pair of coordinates equally across the nine resulting combinations. This additional step was introduced due to our lower confidence in the accuracy of 5′ and 3′ mapped coordinates based on our inability to trim many reads based on their expected sequence composition. This process increases the likelihood of isolating reciprocal pairs of SPO11 oligos (which otherwise requires a precise 2 bp offset between 5′ and 3′ ends of the reciprocal pair), at the expense of 9-fold lowered absolute frequency of any given molecular coordinates.

### Reciprocal filtering of Spo11-oligo libraries to isolate Spo11-DC compatible molecules

To enrich for putative Spo11-DC molecules in both *S. cerevisiae* and mouse libraries, Spo11-oligos were discarded unless a reciprocal partner mapped with 2 bp offset was present within the library. Spo11-DCs were then reported as twice the minimum frequency of either the Franklin or Crick oligo (whichever was lower). Additionally, for quantitative analyses and the plotting of arc-diagrams, all oligos <31 nt were discarded, justified by our physical gel-based analysis of Spo11-DCs in *sae2*Δ cells, and the periodic enrichment of Spo11-DCs for oligos >30 nt.

### Calculating DNA sequence composition at 5′ DSB ends

DNA sequence orientated 5′→3′ around 5′ cleavage sites for the relevant library fraction was aggregated using seqBias (Perl, v5.22.1; https://github.com/Neale-Lab) and plotted as fractional base composition at each base for the top strand.

### Calculating strand disparities

When Spo11 5′ ends are mapped to a reference genome, the 2 bp overhang at cleavage sites translates to a 1 bp offset for the mapped 5′ base on each strand. Thus to compute the relative strand disparity (ratio of the frequency of mapping, in hits per million, HpM, on each strand at a given cleavage site), the mapped Crick coordinates were first shifted by −1 bp. Before calculating disparity ratios, 0.01 HpM were added to both denominator and numerator to avoid errors arising when one of the values was zero. Net, left–right, disparity for individual Spo11-DC molecules was defined as: (Franklin_left_ / Crick_left_) / (Franklin_right_ / Crick_right_).

### Analysis of 6:2 recombination patterns in *msh2*Δ hybrid octad data

All analysed meioses included the *msh2*Δ, which abrogates mismatch repair, thereby enabling the retention of heteroduplex DNA (hDNA) strand information, which then segregates (becoming homoduplex) in the first post-meiotic cell division (octad stage). Whole-genome DNA libraries for each haploid member of every octad were independently prepared and barcoded using Illumina NexteraXT according to manufacturer’s instructions, and sequenced at ∼16-plex on a MiSeq using 2 x 300 bp paired-end reads, obtaining a minimum of at least 20x average genome coverage. Paired-end mapping, genotyping of the ∼65,000 SNP and Indels present in each of the eight haplotypes in each octad, and HR event calling across each octad were performed as described^37^ using publically available scripts (https://github.com/Neale-Lab/OctadRecombinationMapping), generating Event Tables listing position, type, and detailed information describing each isolated HR event (Supplementary Tables XXX). An integral stage of event calling is the partition of the octad into genomic segments of identical adjacent marker segregation (e.g. 5:3, 6:2, 4:4, etc). As with prior work^23^, an inter-event merging threshold of 1.5 kb was used—that is, a minimum of at least 1.5 kb of 4:4 marker segregation was necessary between two adjacent regions of non-4:4 marker segregation for the region to be recorded as two independent HR events. In HR events from *msh2*Δ mutants, segments of 6:2 marker segregation are not expected from simple models of DSB repair, but may arise due to either secondary nicking by an associated nuclease^23^, or from initiation by an adjacent pair of Spo11-DSBs^22^—referred to here as a Spo11-DC.

In order to focus on events that may have arisen from putative gap repair—and compatible with the range of Spo11-DC sizes detected physically (**Fig. 1**)—events were filtered to include only those containing a 6:2 segment, and then further categorised depending upon the estimated length of the 6:2 segment, whether there was more than one 6:2 segment within the event (single vs multi), and the marker segregation patterns flanking the 6:2 segment (categories A, B, C; **Fig. 4** and **Fig. S7**). Because of the perceived length (and visibility) of any segment is affected by local variant density, categorisations are inherently uncertain, but represent our reasonable estimates. Firstly, 6:2 segments were excluded from consideration when the maximum possible length of a 6:2 segment was <30 bp, and when the minimum length of a 6:2 segment was >150 bp. Whilst we don’t exclude the possibility that coincident Spo11-DSBs separated by more than 150 bp might arise (e.g. **Fig. 1c** and **Fig. S1a**; ^18^), we favour the view that the more separated DSBs are, the more likely they will behave as two independent DSBs—each with two recombinationally-active DNA ends—and thus less likely to generate products compatible with a simple model of gap repair. Additionally, most events were classified as Category C (ambiguous) due to lack of useful flanking heteroduplex information. Nevertheless, when possible, flanking heteroduplex segregation patterns were used to exclude some events (Category B, incompatible)—for example, when the flanking hDNA was in *cis*, rather than *trans* orientation (**Fig. 4c**). In other instances strand polarity information—inferred from the overall phasing of the haplotypes based on NCO *trans* hDNA patterns present elsewhere in the octad^23^—were used to aid classification. NCOs without phasing information were classified as Category C (ambiguous). Despite these uncertainties, the fact that the fraction of events falling into Category A is unaffected by deletion of *EXO1, MLH1* or *MLH3* (which when mutated abrogate the generation of 6:2 events arising from spurious nicking^23^), whereas Category B and C are reduced ∼3-fold and ∼7-fold, respectively, by these mutations, provides confidence in the validity of our classifications. Precise categorisation rules are presented below:

NCO events: NCO 6:2 segments flanked by hDNA tracts in *trans* orientation (e.g. CN4; **Fig. S7g**) are considered highly compatible because the hDNA segments are suggestive of repair synthesis tracts. NCO 6:2 segments flanked by hDNA in *cis* orientation are classed as incompatible. NCO 6:2 segment with hDNA on only one side are classified as compatible if the hDNA pattern matches the known phasing of the strands (e.g. CN3; **Fig. S7f**). In this latter case, we infer that the missing information on one side of the event may be absent due to lack of variant coverage. One-sided NCO events with incorrect strand phasing are classified as incompatible. One-sided NCO events where the strand orientation could not be phased were classed as ambiguous.

CO events: There are two places a 6:2 segment can appear; in the centre of the CO exchange (e.g. CC1-CC4; **Fig. S7b-c**) or offset from the CO exchange (e.g. CC5,CC7; **Fig. S7d-e**), but involves one of the two chromatids already involved in the CO, and falls within the 1.5 kb inter-event threshold. Central CO gaps are considered compatible if there are *trans* hDNA patterns either side of the 6:2 segment. However, unlike the case with NCOs, the *trans* patterns can be across two chromatids if the 6:2 segment is at the CO point (e.g. CC1; **Fig. S7b**). In this latter case then the strand orientation phasing can be taken into account, and for all COs where the phasing of both chromatids was known, the hDNA patterns are confirmed to be in *trans* (e.g. CC5; **Fig. S7d**). CO 6:2 segments where strand orientation phasing of either chromatid is not known but display a *trans* hDNA-like pattern are retained in the compatible category (e.g. CC1-2; **Fig. S7b**). COs containing offset 6:2 segments are classed as compatible when the gap has full flanking *trans* hDNA (e.g. CC5; **Fig. S7d**), or half-hDNA which is in the correct phased orientation (e.g. CC7; **Fig. S7e**). COs containing offset 6:2 segments with hDNA in the incorrect orientation (where phasing was possible) are classified as incompatible. Note that the analysis included all CO events, including complex ones showing bi-directional conversions that involve at least two initiating DNA lesions on two non-sister chromatids (e.g. CC3, CC5, CC7; **Fig. S7c-e**)^23^. Such events were included when the outcomes of the initiating lesions could be reasonably anticipated.

### Overlap of 6:2 segments with annotated hotspots

To calculate hotspot overlap, 6:2 segments, up to but not including the non-6:2 flanking markers, were tested for their intersection with the coordinates of a list of previously annotated Spo11-DSB hotspots^16^, generating a binary, yes/no result for each 6:2 segment within each category (A–C). Proportions were then calculated and reported. For these analyses the Spo11-oligo datasets utilised were obtained from SK1 nonhybrid diploids.

### Measuring Spo11 activity within, and surrounding 6:2 segments

To assess the correlation between 6:2 segment locations and local, population average, Spo11-DSB activity, each HR event was partitioned into 6:2 and non-6:2 segments, and the observed amount of Spo11-oligo signal^16^ falling within each partition calculated. For this analysis, 6:2 segments were defined as the region up to but not including the non-6:2 flanking markers. Expected Spo11 signal for the 6:2 segment was calculated based on the fraction of total Spo11-oligo signal expected to fall within this segment were Spo11-DSBs arising uniformly across the entire event region. Finally, observed/expected ratios were calculated for each 6:2 segment within each category (A–C), and plotted as individual points on a log2 scale. Box-and whisker plots indicate median (horizontal bar), upper and lower quartiles of the range (box), and minimum and maximum points within 1.5-fold of interquartile range (whiskers). For these analyses the Spo11-oligo datasets utilised were obtained from SK1 nonhybrid diploids.

### Analysis of *exo1*Δ, *mlh1*Δ and *mlh3*Δ data in *msh2*Δ background

To determine the impact of abrogating the putative nicking activity promoted by Exo1, Mlh1 and Mlh3, previously published datasets^23^ were analysed using the methods described above. Due to the limited number of repeats (2 meioses for each genotype), and the expected similarity in phenotype of each null mutation^23^, these data were pooled and analysed in aggregate (“*exo-mlh*Δ”) in order to increase statistical power.

### Method for handling incomplete octads

Occasionally we encountered octads where a mother-daughter pair had not been separated correctly, resulting in a ‘septad’—identified by a chromatid displaying no visible hDNA information. In these cases, we removed the affected chromatids from our analysis, and thus any HR events arising on these chromatids were subtracted from the total event count.

### Electrophoretic mobility shift assays

DNA substrates were generated by annealing complementary oligos (sequences below). Oligos were mixed in equimolar concentrations (10 mM) in STE (100 mM NaCl, 10 mM Tris-HCl pH 8, 1 mM EDTA), heated and slowly cooled. Substrates were 5′ end-labeled with gamma-^32^P-ATP and T4 polynucleotide kinase and purified by native polyacrylamide gel electrophoresis. The core complex containing Spo11, Rec102, Rec104 and Ski8 from *S. cerevisiae* was purified from baculovirus-infected insect cells (Claeys Bouuaert et al., in preparation).

Binding reactions (20 µl) were carried out in 25 mM Tris-HCl pH 7.5, 7.5% glycerol, 100 mM NaCl, 2 mM DTT, 5 mM MgCl_2_ and 1 mg/ml BSA with 0.5 nM DNA and the indicated concentration of core complexes. Complexes were assembled for 30 minutes at 30 °C and separated on a 5% Tris-acetate-polyacrylamide/bis (80:1) gel containing 0.5 mM MgCl_2_ at 200 V for two hours. Gels were dried, exposed to autoradiography plates and revealed by phosphorimaging.

### Oligonucleotides for substrate preparation

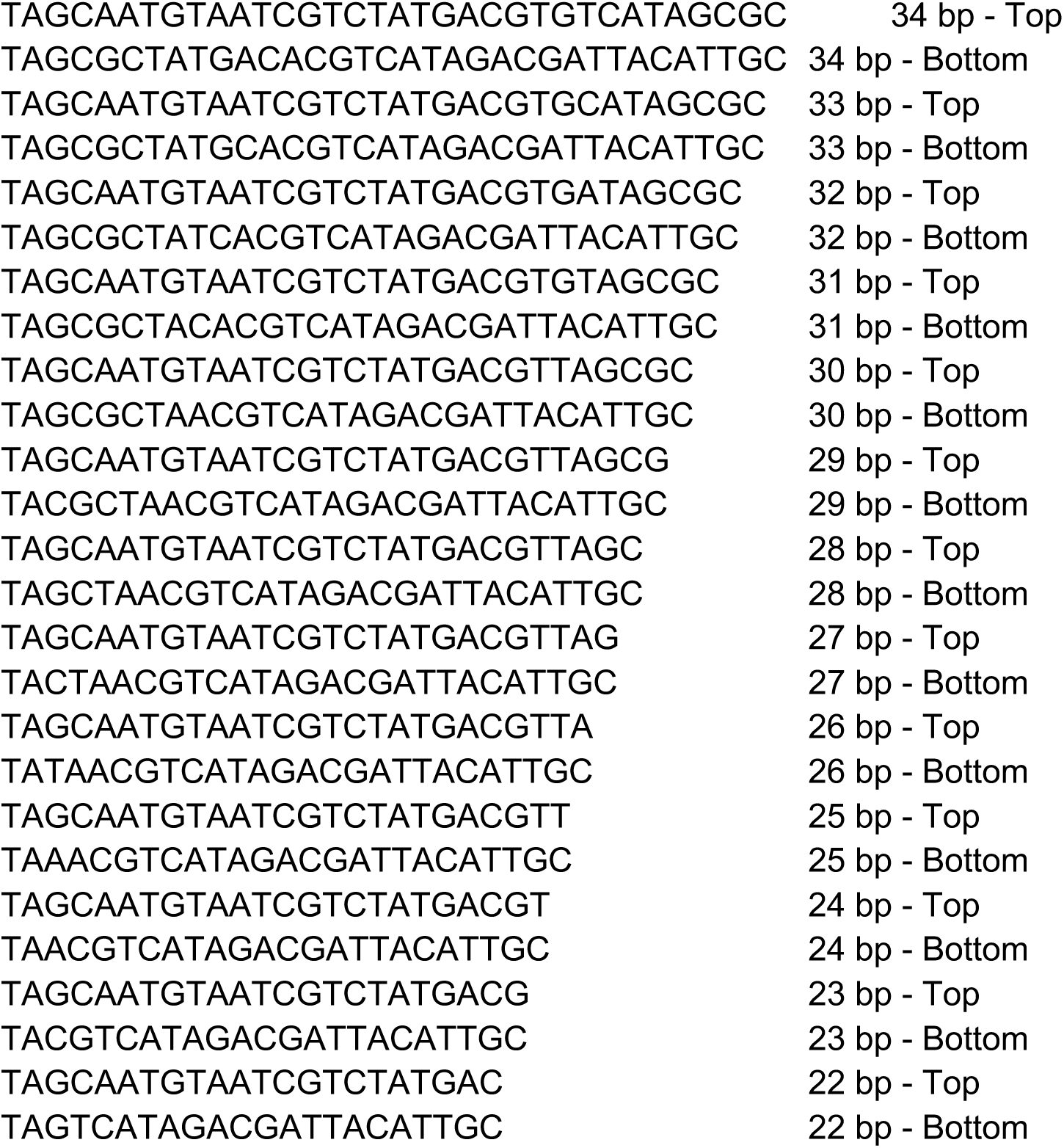

## Supplementary Figures and Legends

**Supplementary Figure 1.**
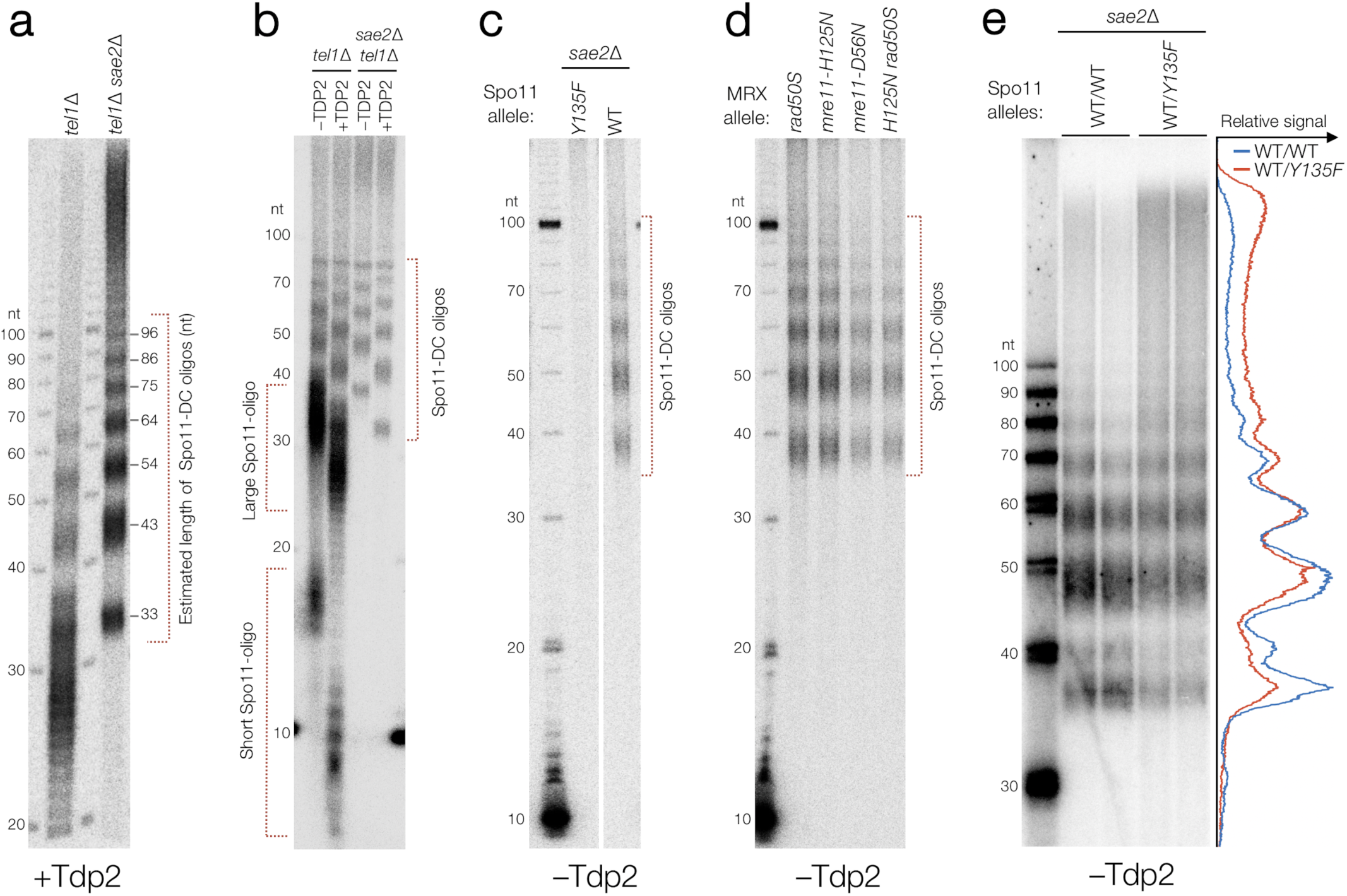
Spo11-DC formation is independent of Mre11-dependent endonucleolytic processing. (Related to Fig. 1) **a-e**, Immunoprecipitated Spo11 oligos and Spo11-DCs isolated from meiotic extracts of the indicated mutants (5 hours after induction of meiosis) were radiolabeled with 3′-dATP using terminal transferase and separated on 19% denaturing PAGE following digestion with proteinase K. In (**a,b**), where indicated, samples are also treated with mammalian TDP2, which removes the residual Spo11 peptide that is left after proteinase K treatment, thereby permitting accurate estimation of the Spo11-DC oligo length^38^. In the absence of TDP2 digestion, the residual 5′-linked Spo11 peptide retards migration of Spo11 oligos and Spo11-DCs by the equivalent of ∼6-8 nt (**b-e**). A 10-nt ladder (also radiolabelled with 3′-dATP) is included in each gel to permit accurate sizing. As expected, Spo11-DCs are abolished by homozygous mutation of the Spo11 active site (*spo11-Y135F), (***c**), and arise independently of single or dual mutations in the MRX complex (*rad50S, mre11-H125N, mre11-D56N*) that abrogate endonuclease activity^8,20^ (**d**). *SPO11*/*spo11-Y135F* heterozygous diploids display an altered Spo11-DC oligo size distribution (biological duplicate lanes of each are presented alongside averaged intensity trace) (**e**).

**Supplementary Figure 2.**
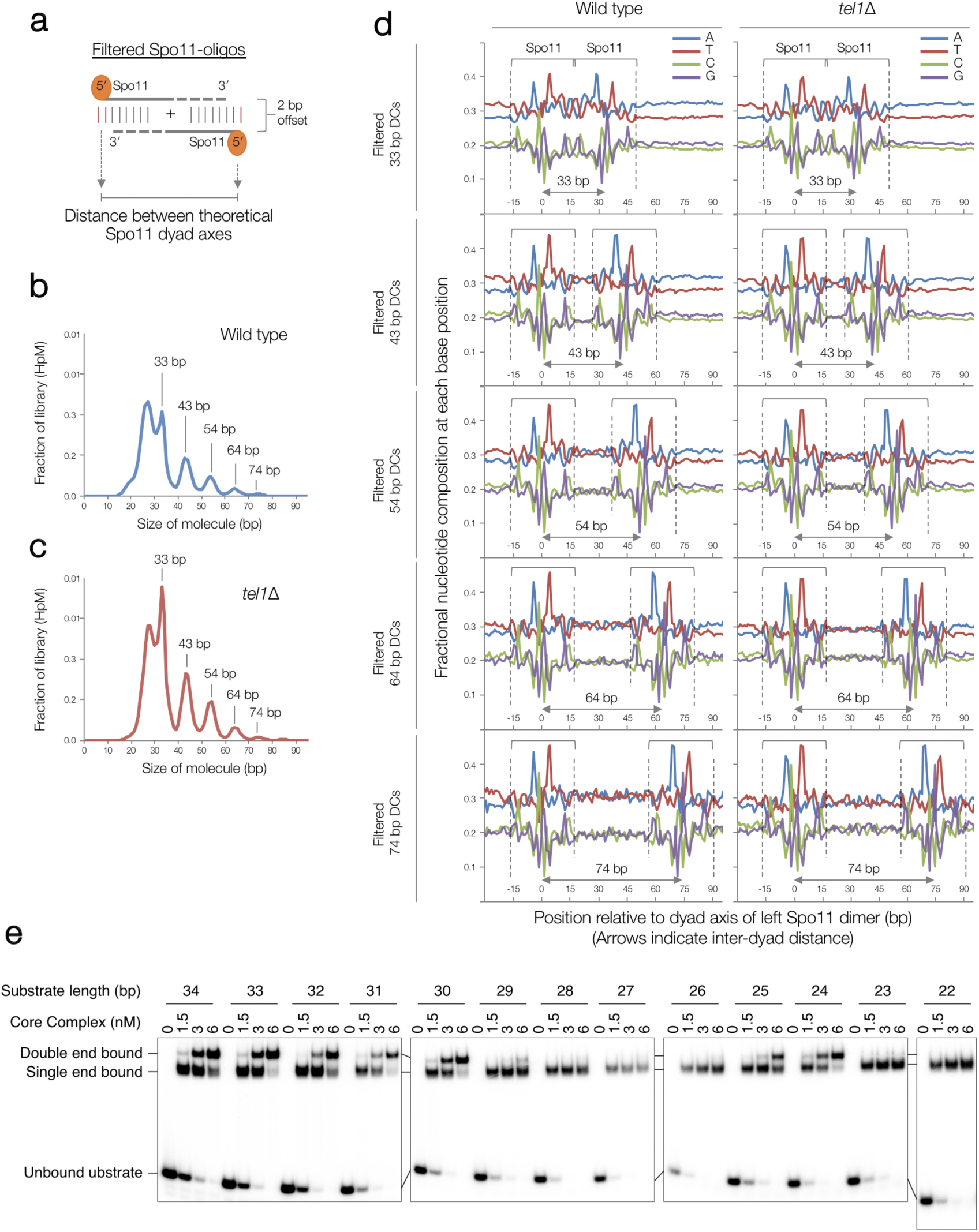
Biased sequence composition around Spo11-DC 5′ ends. (Related to Fig. 2) **a**, Cartoon showing how Spo11-DC were filtered out from total Spo11-oligo libraries based on overlapping molecules sharing 5′ and 3′ coordinates with a precise 2 bp offset. The theoretical dyad axis of each Spo11 dimer (at each end of the molecule) is indicated. Due to the rotational symmetry of cleavage, the distance between such dyad axes is identical to the filtered oligo length. **b-c**, Size distribution of filtered Spo11 oligos in wild type and *tel1*Δ strains. Periodic peaks in the distribution are indicated. **d**, The mean nucleotide composition of filtered Spo11 oligos of the indicated size was computed for each base position and plotted relative to the inferred dyad axis of cleavage of the leftmost Spo11 DSB, revealing signature nucleotide skews characteristic of Spo11 at both the 5′ and 3′ ends. No base skews were observed in the central regions of each molecule, arguing against a major influencer of Spo11-DC formation being localised DNA bending, which is expected to be favoured by an AT-rich base composition. **e**, *In vitro* DNA mobility shift assay. Spo11 core complex (Spo11, Rec102, Rec104, Ski8) was incubated with double-strand DNA substrate of different lengths. Quantification is provided in **Fig. 2**j.

**Supplementary Figure 3.**
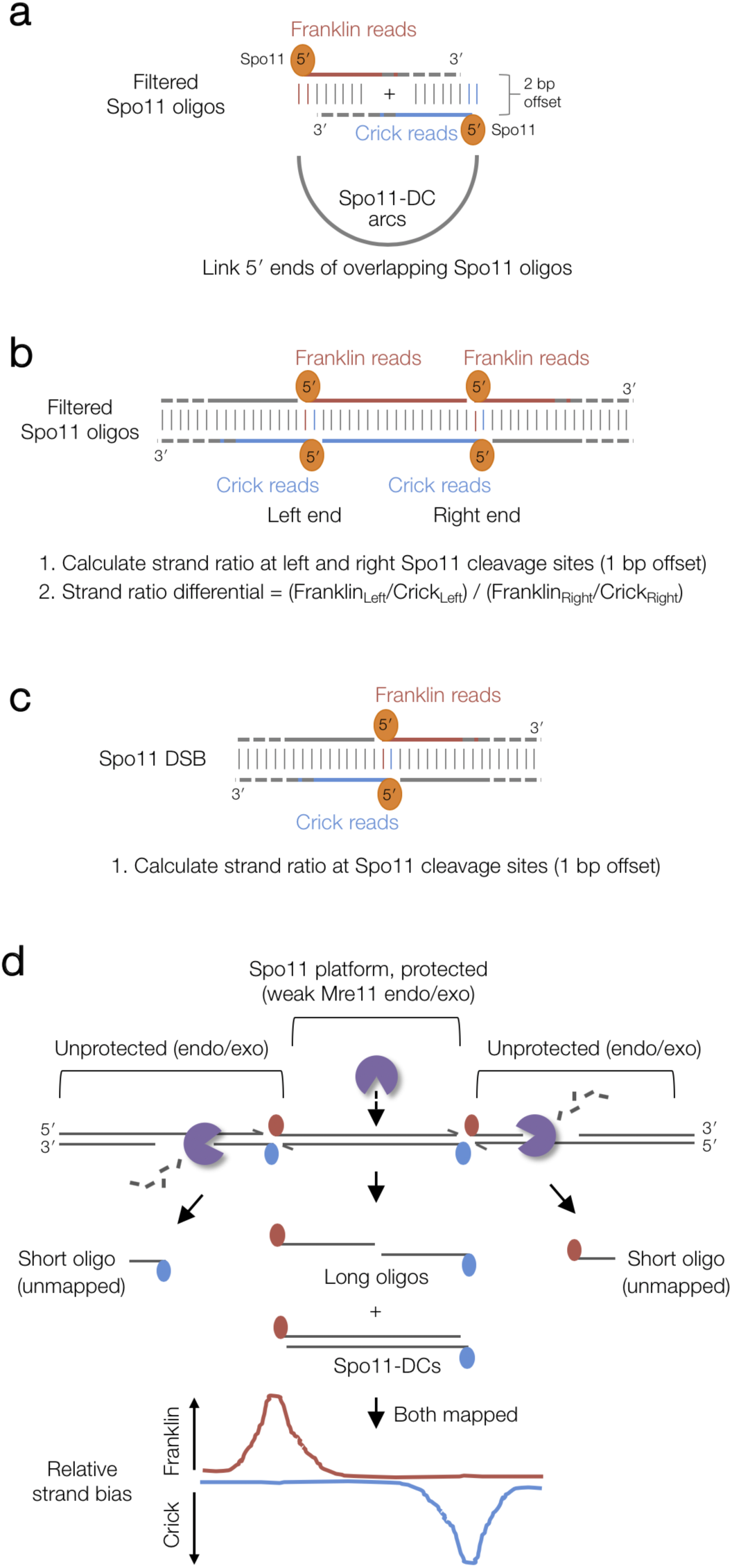
Cartoon to explain the mapped strand disparity. (Related to Fig. 3 and Fig. 6) **a**, Spo11-DC arcs link the 5′ ends of overlapping Franklin- and Crick-strand filtered reads. **b**, Explanatory cartoon for the calculation of Spo11-DC strand ratio and strand-ratio differential (Left/Right). **c**, Strand-ratio calculation for all Spo11 oligos. **d**, Model to account for observed strand bias. Mre11-dependent 3′→5′ exonuclease activity is shown relative to the covalent attachment of Spo11 to 5′ DNA ends. Spo11 oligos protected from Mre11 exonuclease activity within the axis-associated Spo11 platform are efficiently mapped (long Spo11 oligos), while efficient resection in the flanking regions leads to shorter Spo11 oligos that are not mapped. DNA bound by Spo11 platform may be less efficiently cleaved by Mre11 endonuclease (dashed arrow) because of steric constraints and/or limited access between the Spo11 dimers.

**Supplementary Figure 4.**
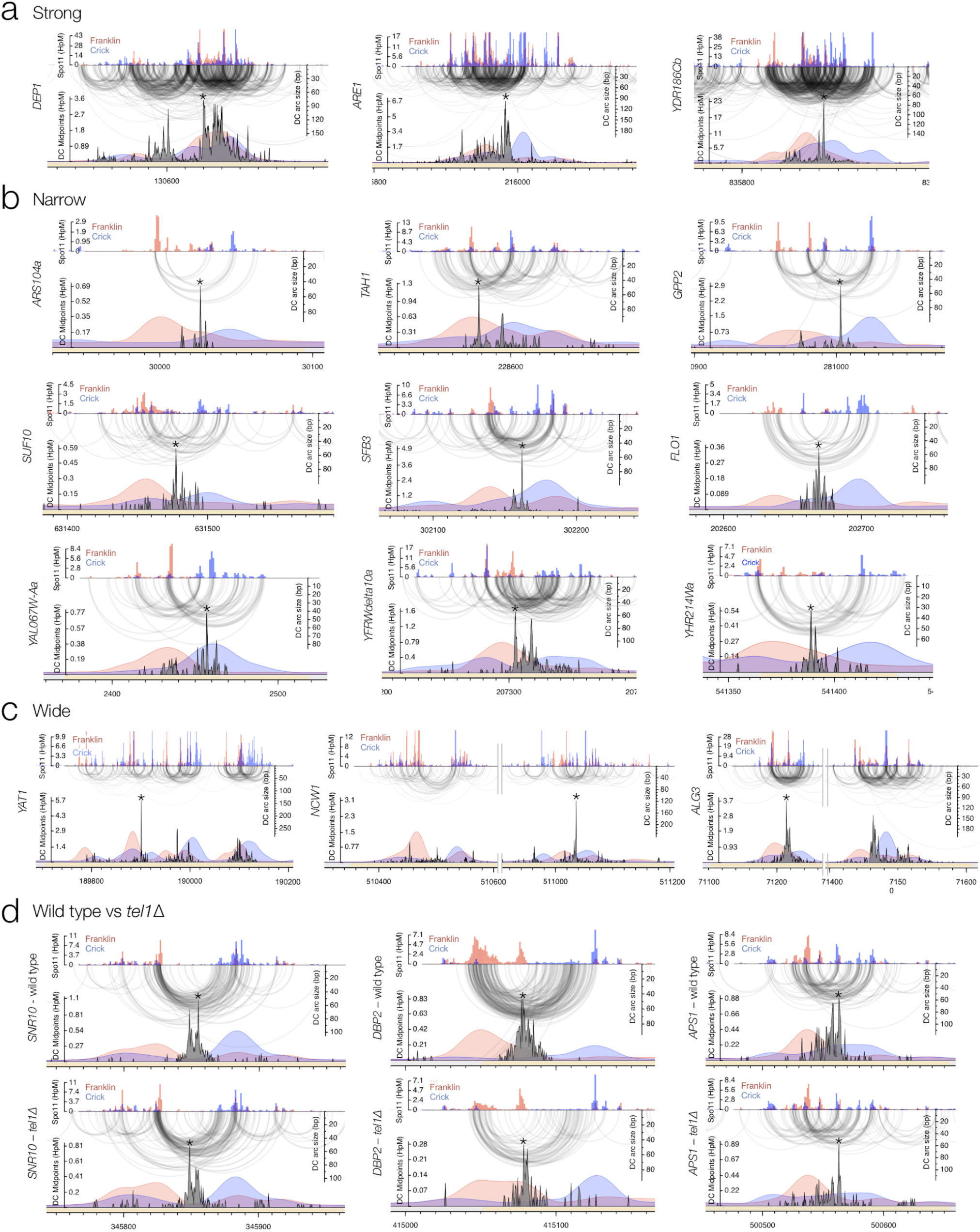
Fine-scale analysis of Spo11-DCs within representative classes of DSB hotspots. (Related to Fig. 3) **a-d**, Arc diagram depiction of Spo11-DCs mapped across representative hotspots encompassing strong (**a**), narrow (**b**), and wide (**c**) classes, presented as in **Fig. 3**. In (**d**), wild-type and *tel1*Δ data are compared for the same hotspots.

**Supplementary Figure 5.**
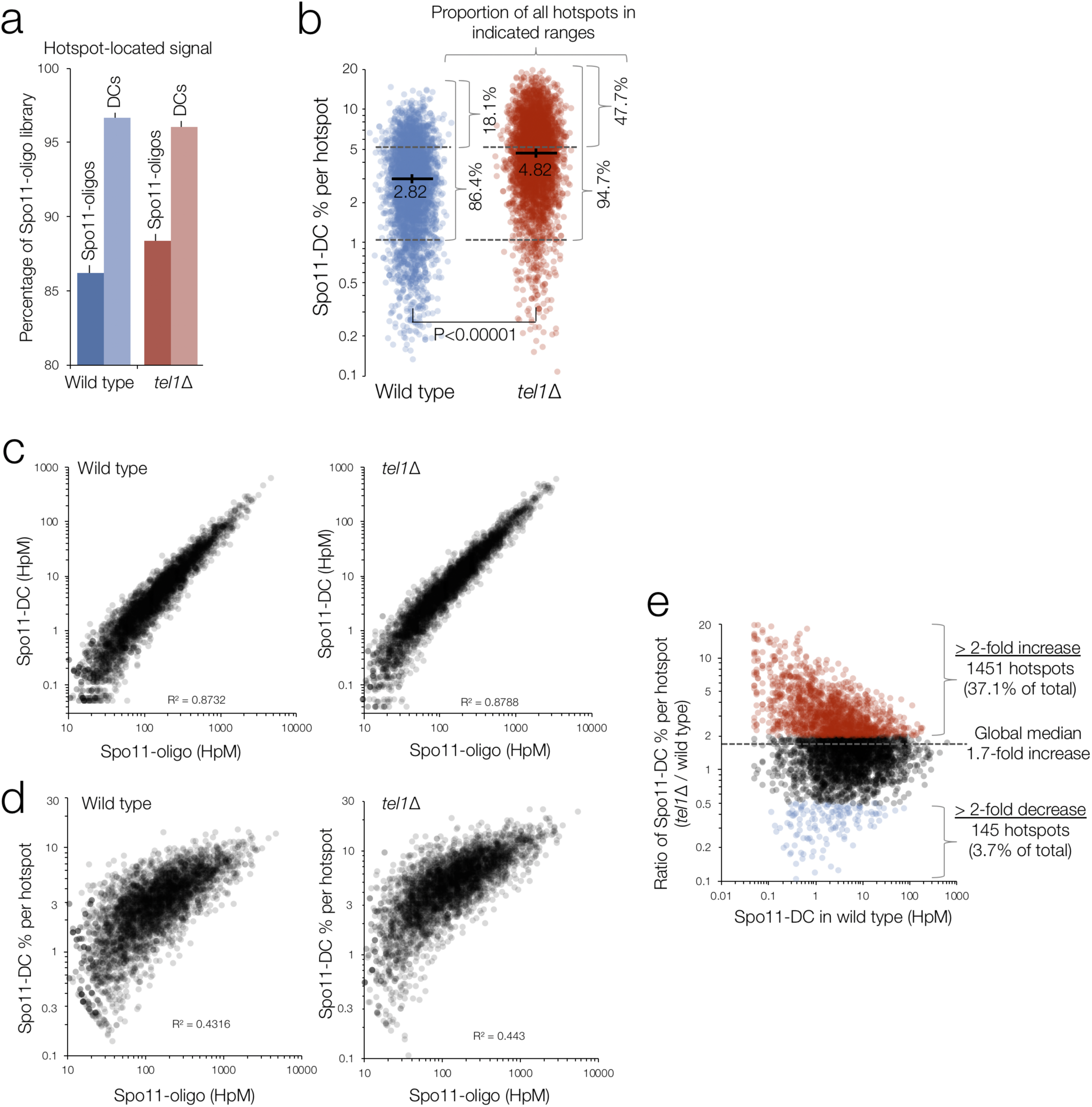
Analysis of Spo11-DC composition of DSB hotspots in wild type and *tel1*Δ. (Related to Fig. 3) **a**, Percentage of total Spo11 oligos and filtered Spo11-DCs that arise within annotated DSB hotspots. **b**, Percentage of total Spo11 oligos that are Spo11-DCs, plotted for every hotspot. P value indicates Kruskal-Wallis H-test. **c-d**, Quantitative correlation between filtered Spo11-DC frequency (**c**), or percentage of Spo11-DCs within each hotspot (**d**), and total Spo11-oligo frequency for all DSB hotspots **e**, Comparison between *tel1*Δ and wild type of the percentage of total Spo11-oligo signal within each hotspot that is classified as a Spo11-DC. These ratios are stratified on the X-axis by the Spo11-DC frequency in wild type cells, and ratios are coloured to indicate those hotspots where the proportion of Spo11-DCs is at least 2-fold increased (red) or 2-fold decreased (blue) in *tel1*Δ relative to wild type.

**Supplementary Figure 6.**
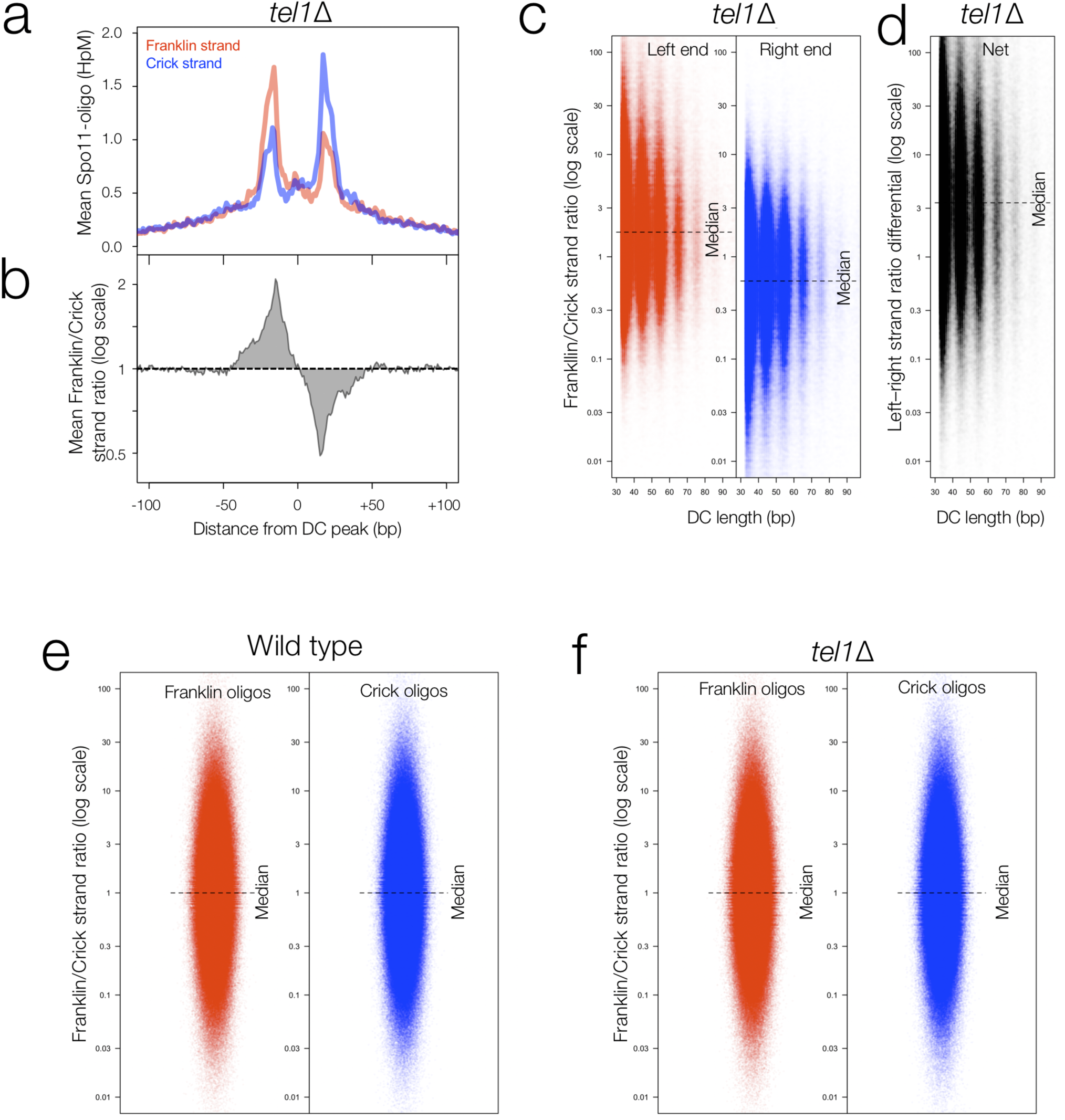
Global analysis of strand disparities at Spo11-DC termini in *tel1*Δ cells. (Related to Fig. 3) **a-e**, Strand ratio of Spo11 oligos at Spo11-DC sites is a genome-wide feature also in *tel1*Δ cells. Average strand-specific Spo11-oligo signal (**a**), and strand ratio (**b**), centred upon the strongest Spo11-DC midpoint within every DSB hotspot. Strand ratio (Franklin/Crick total Spo11-oligo HpM) was computed at the left and right 5′ end of every unique Spo11-DC molecule (**Fig. S3b**), stratified by length (**c**). Strand-ratio differential (**d**) indicates the fold difference in the ratios when comparing the left and right 5′ ends of each Spo11-DC molecule (**Fig. S3b**). **e-f**, Strand ratio (Franklin/Crick total Spo11-oligo HpM) was computed at the 5′ end of all observed (unfiltered) Franklin or Crick strand Spo11 oligos (as in **Fig. S3c**). Unlike at Spo11-DC sites (**Fig. 3f-h**), bulk Spo11 oligos, across all sites, display no strand disparity in either wild type (**e**) or *tel1*Δ (**f**) strains.

**Supplementary Figure 7.**
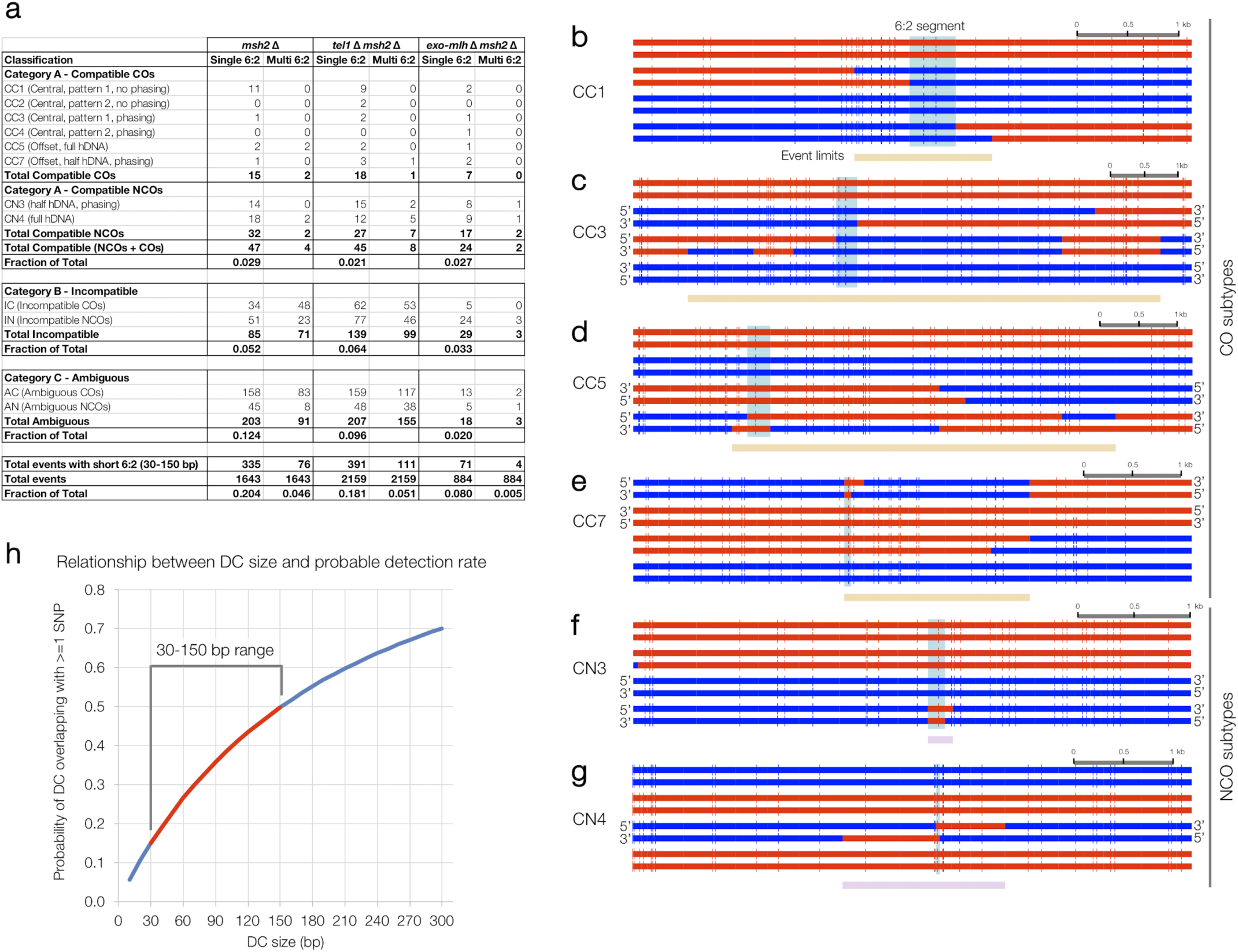
Categorisation of events containing short 6:2 segments. (Related to Fig. 4) **a**, Summary of frequencies of each subclassification event type for the indicated strains. Only events containing 6:2 segments 30 to 150 bp in length are considered. Events were separated into those with a single or multiple such 6:2 segments. Fractions of total for each subtype are not calculated for multi events because they frequently contain more than one sub-type. **b-g**, Example event sub-classifications. Genotype calls are made at each marker (vertical line). Adjacent segments of the same genotype are joined with horizontal bars (red or blue) to aid visualisation of patterns. Each horizontal bar is sequenced haplotype from one meiotic octad. 6:2 segments are indicated in pale blue. Event limits are indicated by beige (crossover) or pink (noncrossover) bars. Orientation of 5′ and 3′ strands are indicated in instances where it was possible to obtain phasing information from noncrossover *trans* events within the event, or from events elsewhere in the octad. In (**c**), a second segment of 6:2 segregation is not considered because the minimum length is >1.2 kb. **h**, To estimate probable detection rates of theoretical Spo11-DC of varying size, sliding windows of increasing size were moved across the reference genome, and the number of genetic markers within each window was recorded for each position. As examples, on average, Spo11-DCs 30 bp and 150 bp in size will be detected only 15% and 50% of the time, respectively. We note that due to the non-uniform distribution of genetic markers—in particular the slightly greater density within intergenic regions where Spo11 DSBs most often arise—the probability of detection may be slightly greater than that estimated from the genome-wide polymorphism density.

**Supplementary Figure 8.**
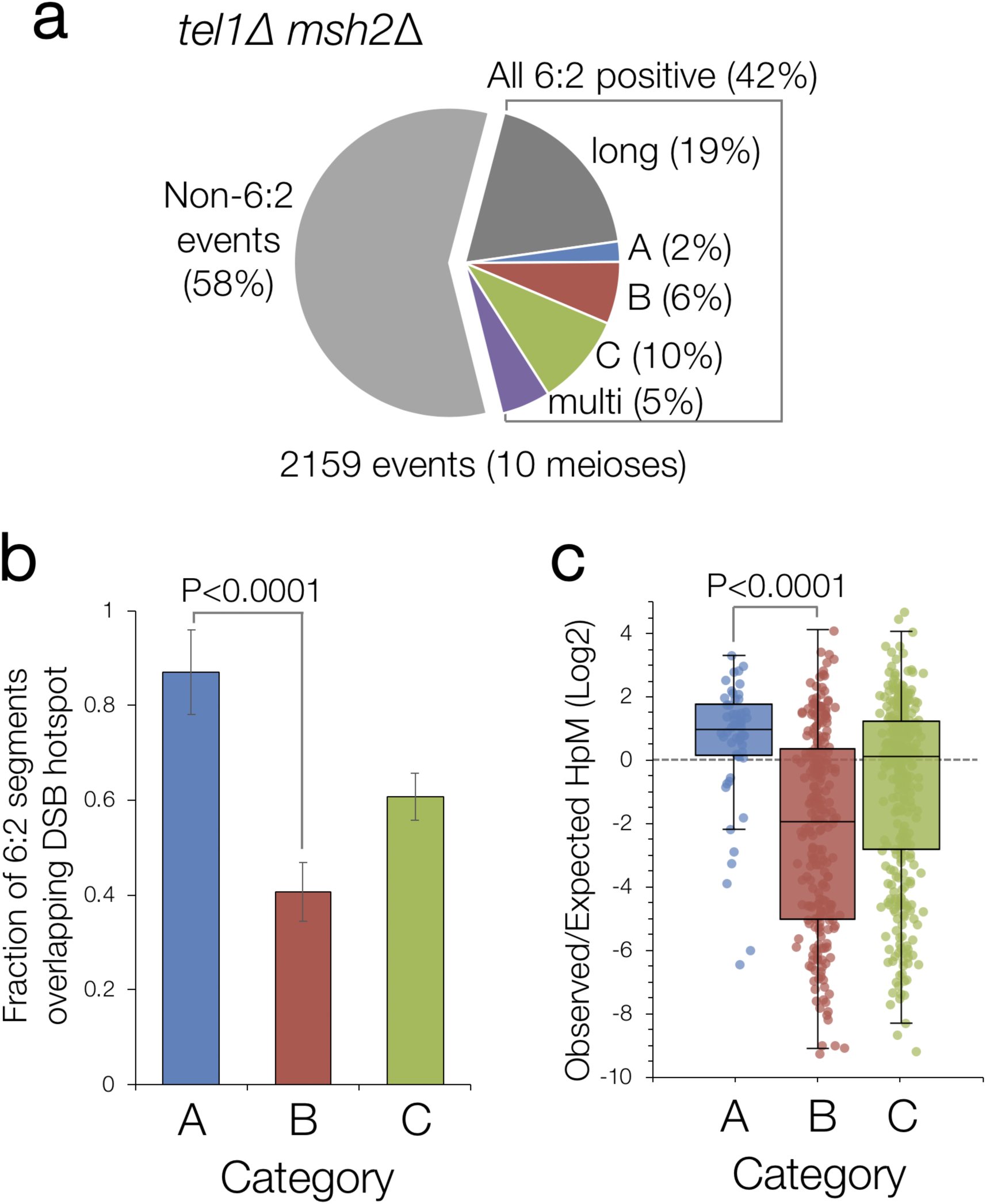
Identification of gap repair events during meiosis in *tel1*Δ cells. (Related to Fig. 4) **a**, Quantification of recombination event types in *tel1*Δ *msh2*Δ based on categories presented in **Fig. 4c. b**, Fraction of 6:2 segments (*tel1*Δ *msh2*Δ) overlapping hotspots. **c**, Log_2_ ratio of observed Spo11-oligo density (using *tel1*Δ Spo11-oligo data^16^ within each 6:2 segment divided by the mean Spo11-oligo density within the entire event, in *tel1*Δ *msh2*Δ, for each category. P values indicate Z-test of proportions (**b**) and Kruskal-Wallis H-test (**c**).

**Supplementary Figure 9.**
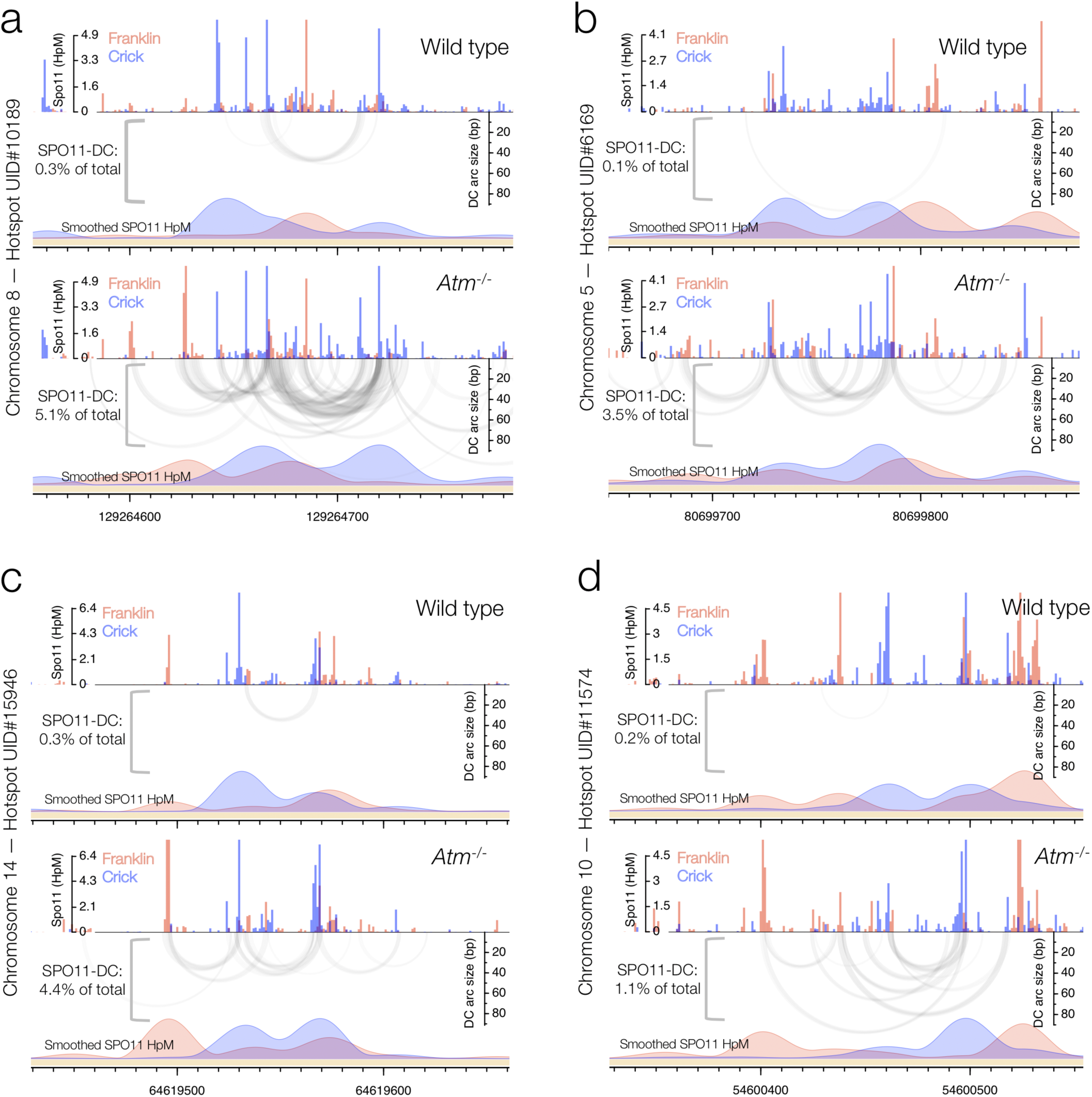
Fine-scale analysis of SPO11-DCs within representative mouse DSB hotspots in wild type and *Atm*^*-/-*^. (Related to Fig. 5) **a-d**, Arc diagrams of SPO11-DCs (grey-scale frequency-weighted arcs) in wild-type and *Atm*^*-/-*^ relative to total strand-specific SPO11 oligos (upper, raw; lower, smoothed; red, Franklin; blue, Crick). Percentage of total SPO11 oligos that are SPO11-DCs is indicated.

**Table S1.**
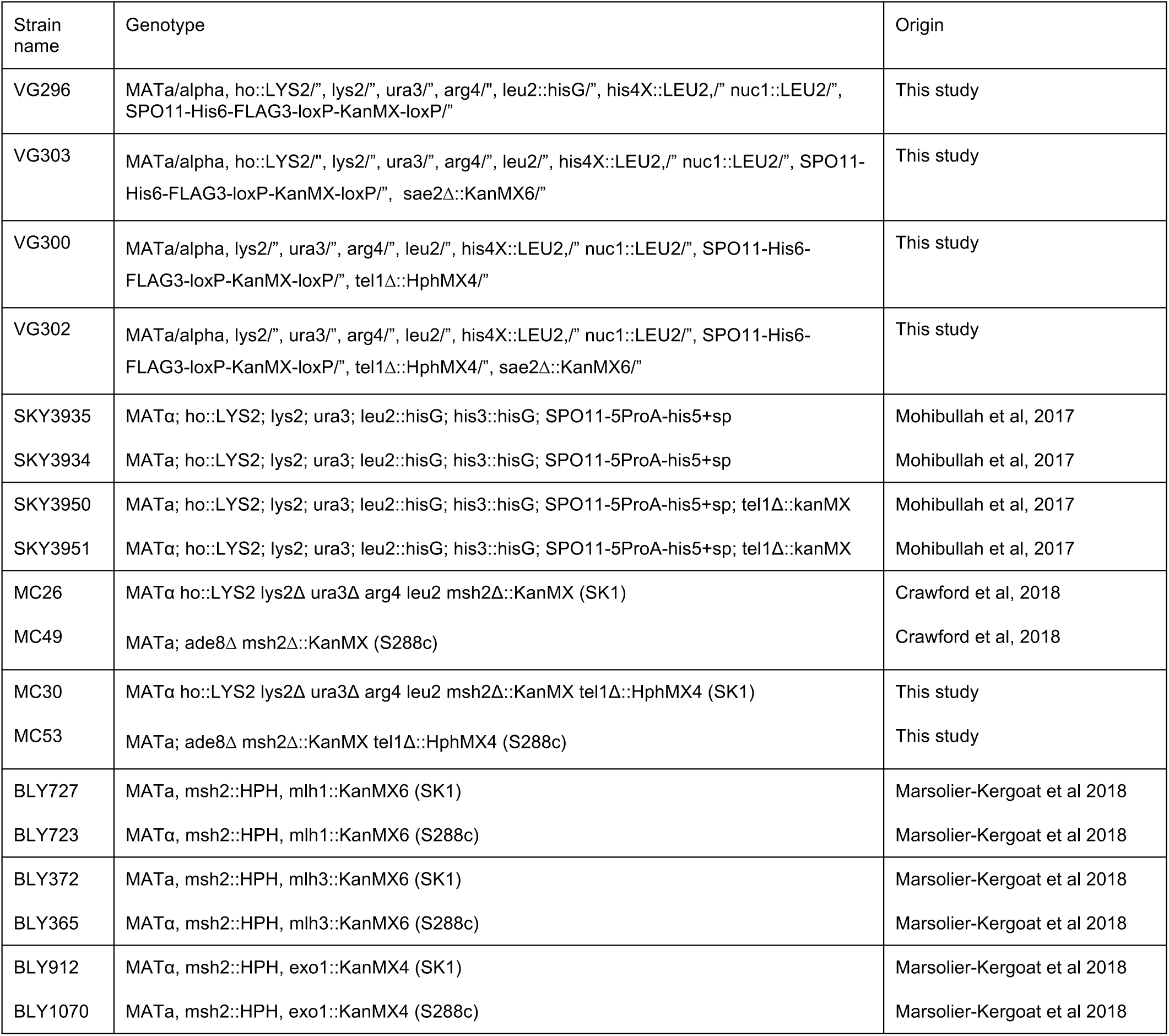
Strains used in this study. Unless indicated, all strains are of the SK1 genetic background^40^. For recombination mapping in octads, haploids were mated directly prior to sporulation as described^23,37^.

## References

1. Szostak, J. W., Orr-Weaver, T. L., Rothstein, R. J. & Stahl, F. W. The double-strand-break repair model for recombination. Cell 33, 25–35 (1983).

2. Keeney, S., Giroux, C. N. & Kleckner, N. Meiosis-specific DNA double-strand breaks are catalyzed by Spo11, a member of a widely conserved protein family. Cell 88, 375–384 (1997).

3. Bergerat, A. et al. An atypical topoisomerase II from Archaea with implications for meiotic recombination. Nature 386, 414–417 (1997).

4. Liu, J., Wu, T. C. & Lichten, M. The location and structure of double-strand DNA breaks induced during yeast meiosis: evidence for a covalently linked DNA-protein intermediate. EMBO J 14, 4599–4608 (1995).

5. de Massy, B., Rocco, V. & Nicolas, A. The nucleotide mapping of DNA double-strand breaks at the CYS3 initiation site of meiotic recombination in Saccharomyces cerevisiae. EMBO J 14, 4589–4598 (1995).

6. Keeney, S. & Kleckner, N. Covalent protein-DNA complexes at the 5’ strand termini of meiosisspecific double-strand breaks in yeast. Proc Natl Acad Sci U S A 92, 11274–11278 (1995).

7. Furuse, M. et al. Distinct roles of two separable in vitro activities of yeast Mre11 in mitotic and meiotic recombination. EMBO J 17, 6412–6425 (1998).

8. Moreau, S., Ferguson, J. R. & Symington, L. S. The nuclease activity of Mre11 is required for meiosis but not for mating type switching, end joining, or telomere maintenance. Mol Cell Biol 19, 556–566 (1999).

9. Neale, M. J., Pan, J. & Keeney, S. Endonucleolytic processing of covalent protein-linked DNA double-strand breaks. Nature 436, 1053–1057 (2005).

10. Garcia, V., Phelps, S. E. L., Gray, S. & Neale, M. J. Bidirectional resection of DNA doublestrand breaks by Mre11 and Exo1. Nature 479, 241–244 (2011).

11. Pan, J. et al. A Hierarchical Combination of Factors Shapes the Genome-wide Topography of Yeast Meiotic Recombination Initiation. Cell 144, 719–731 (2011).

12. Fowler, K. R., Sasaki, M., Milman, N., Keeney, S. & Smith, G. R. Evolutionarily diverse determinants of meiotic DNA break and recombination landscapes across the genome. Genome Res 24, 1650–1664 (2014).

13. Lam, I. & Keeney, S. Nonparadoxical evolutionary stability of the recombination initiation landscape in yeast. Science 350, 932–937 (2015).

14. Choi, K. et al. Nucleosomes and DNA methylation shape meiotic DSB frequency inArabidopsis thalianatransposons and gene regulatory regions. Genome Res (2018).

15. Lange, J. et al. The Landscape of Mouse Meiotic Double-Strand Break Formation, Processing, and Repair. Cell (2016).

16. Mohibullah, N. & Keeney, S. Numerical and spatial patterning of yeast meiotic DNA breaks by Tel1. Genome Res 27, 278–288 (2017).

17. Lange, J. et al. ATM controls meiotic double-strand-break formation. Nature 479, 237–240 (2011).

18. Garcia, V., Gray, S., Allison, R. M., Cooper, T. J. & Neale, M. J. Tel1(ATM)-mediated interference suppresses clustered meiotic double-strand-break formation. Nature 520, 114–118 (2015).

19. Cannavo, E. et al. Regulatory control of DNA end resection by Sae2 phosphorylation. Nat Commun 9, 4016 (2018).

20. Cao, L., Alani, E. & Kleckner, N. A pathway for generation and processing of double-strand breaks during meiotic recombination in S. cerevisiae. Cell 61, 1089–1101 (1990).

21. Gittens, W. et al. A nucleotide resolution map of Top2-linked DNA breaks in the yeast and human genome. bioRxiv (2019).

22. Martini, E. et al. Genome-wide analysis of heteroduplex DNA in mismatch repair-deficient yeast cells reveals novel properties of meiotic recombination pathways. PLoS Genet 7, e1002305 (2011).

23. Marsolier-Kergoat, M. C., Khan, M. M., Schott, J., Zhu, X. & Llorente, B. Mechanistic View and Genetic Control of DNA Recombination during Meiosis. Mol Cell 70, 9-20.e6 (2018).

24. Keeney, S. Spo11 and the Formation of DNA Double-Strand Breaks in Meiosis. Genome dynamics and stability 2, 81 (2008).

25. Diaz, R. L., Alcid, A. D., Berger, J. M. & Keeney, S. Identification of residues in yeast Spo11p critical for meiotic DNA double-strand break formation. Mol Cell Biol 22, 1106–1115 (2002).

26. Noll, M. Internal structure of the chromatin subunlt. Nucleic Acids Research 1, 1573–1578 (1974).

27. Gittens, W. H. et al. A nucleotide resolution map of Top2-linked DNA breaks in the yeast and human genome. Nature Communications 10, (2019).

28. Prieler, S., Penkner, A., Borde, V. & Klein, F. The control of Spo11’s interaction with meiotic recombination hotspots. Genes Dev 19, 255–269 (2005).

29. Panizza, S. et al. Spo11-accessory proteins link double-strand break sites to the chromosome axis in early meiotic recombination. Cell 146, 372–383 (2011).

30. Kugou, K. et al. Rec8 guides canonical Spo11 distribution along yeast meiotic chromosomes. Mol Biol Cell 20, 3064–3076 (2009).

31. Li, J., Hooker, G. W. & Roeder, G. S. Saccharomyces cerevisiae Mer2, Mei4 and Rec114 form a complex required for meiotic double-strand break formation. Genetics 173, 1969–1981 (2006).

32. Kee, K., Protacio, R. U., Arora, C. & Keeney, S. Spatial organization and dynamics of the association of Rec102 and Rec104 with meiotic chromosomes. EMBO J 23, 1815–1824 (2004).

33. Kumar, R., Bourbon, H. M. & de Massy, B. Functional conservation of Mei4 for meiotic DNA double-strand break formation from yeasts to mice. Genes Dev 24, 1266–1280 (2010).

34. Zhang, L., Kleckner, N. E., Storlazzi, A. & Kim, K. P. Meiotic double-strand breaks occur once per pair of (sister) chromatids and, via Mec1/ATR and Tel1/ATM, once per quartet of chromatids. Proc Natl Acad Sci U S A 108, 20036–20041 (2011).

35. Joyce, E. F. et al. Drosophila ATM and ATR have distinct activities in the regulation of meiotic DNA damage and repair. J Cell Biol 195, 359–367 (2011).

36. Carballo, J. A. et al. Budding Yeast ATM/ATR Control Meiotic Double-Strand Break (DSB) Levels by Down-Regulating Rec114, an Essential Component of the DSB-machinery. PLoS Genet 9, e1003545 (2013).

37. Crawford, M., Cooper, T. J., Marsolier-Kergoat, M.-C., Llorente, B. & Neale, M. J. Separable roles of the DNA damage response kinase Mec1(ATR) and its activator Rad24(RAD17) within the regulation of meiotic recombination. bioRxiv (2018).

38. Johnson, D., Allison, R. M., Cannavo, E., Cejka, P. & Neale, M. Removal of Spo11 from meiotic DNA breaks in vitro but not in vivo by Tyrosyl DNA Phosphodiesterase 2. bioRxiv http://dx.doi.org/10.1101/527333, (2019).

39. Xu, L. & Kleckner, N. Sequence non-specific double-strand breaks and interhomolog interactions prior to double-strand break formation at a meiotic recombination hot spot in yeast. EMBO J 14, 5115–5128 (1995).

40. Kane, S. M. & Roth, R. Carbohydrate metabolism during ascospore development in yeast. J Bacteriol 118, 8–14 (1974).

